# Zuranolone mitigates delirium-like bispectral EEG changes, behavioral deficits, and neuroinflammation across surgical and inflammatory mouse models and age groups

**DOI:** 10.64898/2026.05.19.726175

**Authors:** Bun Aoyama, Shota Nishitani, Kyosuke Yamanishi, Hieu D Nguyen, Rika Sakuma, Takaya Ishii, Yukiko Ikeda, Tsuyoshi Nishiguchi, Ilgin Genc, Nathan J Phuong, Nipun Gorantla, Tomoteru Seki, Akiyoshi Shimura, Takashi Kawano, Gen Shinozaki

## Abstract

Delirium is an acute, fluctuating brain dysfunction that commonly follows surgery and systemic inflammation, disproportionately affects older adults, and remains difficult to quantify continuously over time and treat pharmacologically. Here, we tested whether the neuroactive steroid zuranolone, a positive allosteric modulator of synaptic and extrasynaptic GABA_A receptors, mitigates delirium-like abnormalities across two complementary murine delirium models, a lipopolysaccharide-induced systemic inflammation (LPS) model and a postoperative delirium (POD) model, primarily in young and aged mice, with selected analyses in super-aged mice. Using continuous EEG with a validated bispectral EEG (BSEEG) metric, we found that zuranolone attenuated delirium-like EEG slowing in the LPS model in young mice in a dose-dependent manner and retained efficacy in aged mice. In the POD model, prophylactic dosing provided limited benefit in young mice, whereas post-surgery treatment reduced postoperative BSEEG elevations. In aged mice, prophylactic dosing suppressed POD-associated BSEEG abnormalities, and in super-aged mice, prophylactic zuranolone improved survival after POD induction. In parallel, zuranolone reduced microglial density and activation markers (IBA1 and CD68 immunoreactivity) at 24 h after POD surgery and after LPS challenge, with effects that were particularly evident in peri-screw site tissue in young POD mice and more broadly distributed across regions in aged mice. Finally, in young mice, zuranolone improved a composite behavioral severity score in the LPS model, whereas behavioral effects in the POD model were modest and domain-specific. Together, these findings support zuranolone as a candidate strategy to reduce delirium-like electrophysiological and neuroimmune abnormalities, with the strongest effects in inflammation-driven and age-vulnerable contexts.

## Introduction

Delirium is a common and severe form of acute brain dysfunction characterized by fluctuating disturbances in attention, awareness, and cognition. It occurs frequently in hospitalized patients, particularly after surgery and during systemic inflammatory states, and is associated with increased mortality, prolonged hospitalization, long-term cognitive decline, and substantial healthcare cost [1] [2]. Despite long-standing clinical recognition, delirium remains difficult to quantify longitudinally and difficult to treat pharmacologically [3] [4].

Advanced age is the strongest and most consistent risk factor for delirium across clinical contexts [5] [6]. Older adults not only exhibit a higher incidence of delirium but also experience more severe and prolonged courses, with incomplete recovery in some cases [7]. Experimental and clinical evidence suggests that age-related vulnerability reflects interaction among baseline neural fragility, heightened neuroimmune responses, and impaired homeostatic mechanisms—often described as “inflammaging” and microglial priming [8]. These age-related changes may contribute to atypical, subtle, or hypoactive presentations, and together with the fluctuating nature of delirium over time, clinical assessments alone can miss transient episodes and limit accurate tracking of disease severity and recovery, particularly in older adults.

Electroencephalography (EEG) has long been recognized as a sensitive and objective marker of delirium. Across etiologies, delirium is associated with characteristic EEG slowing, increased low-frequency power, and disruption of normal diurnal rhythmicity [9]. We previously developed and validated a bispectral EEG (BSEEG) metric based on the ratio of low-to high-frequency power. In prior clinical and preclinical studies, this measure was shown to be sensitive to delirium-associated brain dysfunction in both humans and mice, supporting its use as a translational EEG-derived method across species [10] [11] [12] [13]. This approach enables continuous, longitudinal assessment of brain dysfunction triggered by inflammatory insults ranging from the acute response to the subsequent recovery phase.

Systemic inflammation is a major precipitant of delirium in both surgical and medical settings. Lipopolysaccharide (LPS) administration in rodents induces a robust inflammatory response accompanied by sickness behavior, cognitive deficits, and EEG slowing recapitulating key features of inflammation-associated delirium [8]. Similarly, surgical trauma triggers peripheral immune activation, blood–brain barrier perturbation, and neuroinflammation [14] [15]. Microglial activation has emerged as a key mechanism of neuroinflammation linking peripheral insults to delirium-like brain dysfunction, with exaggerated and prolonged responses observed in aged brains [6]. However, pharmacologic interventions that attenuate delirium-like electrophysiological and behavioral abnormalities across age groups remain limited.

Zuranolone is an orally bioavailable neuroactive steroid related to allopregnanolone that acts as a positive allosteric modulator of GABA_A receptors [16] [17]. Neuroactive steroids such as allopregnanolone can modulate both synaptic and extrasynaptic GABA_A receptor signaling, thereby influencing phasic and tonic inhibition through mechanisms that are pharmacologically distinct from classical benzodiazepines [16] [18]. This profile may be relevant to delirium because acute systemic inflammation and perioperative stress are associated with disturbances in arousal regulation and cognitive function, processes in which GABA_A receptor–mediated inhibition has an important modulatory role [19] [20] [21]. Rather than serving as a direct anti-inflammatory strategy, zuranolone may modulate downstream brain responses to inflammatory or surgical stress by enhancing GABA_A receptor–mediated inhibitory signaling during vulnerable brain states [16] [21]. Zuranolone was approved by the U.S. Food and Drug Administration in August 2023 for postpartum depression, supporting its clinical feasibility [22] [23]. In addition, phase 2 and phase 3 clinical trials and meta-analyses have evaluated its efficacy in adults with major depressive disorder [24]. Together, these features provide a mechanistic and clinically tractable rationale for testing zuranolone in inflammation- and surgery-associated delirium models.

In the present study, we investigated whether zuranolone mitigates delirium-like electrophysiological, behavioral, and neuroinflammatory abnormalities across two complementary murine models: a postoperative delirium model induced by EEG head-mount implantation surgery (POD model) and a lipopolysaccharide-induced systemic inflammation model (LPS model), across young, aged, and super-aged mice. Using continuous EEG recording with a validated BSEEG metric, we quantified longitudinal electrophysiological changes during predefined acute windows as well as extended recovery periods. We further assessed delirium-relevant behavioral outcomes and microglial activation to link electrophysiological changes with behavioral and cellular correlates. By systematically assessing diverse age groups (young, aged, and super-aged), multiple insult types (POD vs. LPS), and different timings of zuranolone administration (prevention vs. treatment), this study aimed to evaluate the potential therapeutic benefit of zuranolone in attenuating delirium-like brain dysfunction.

## Methods

### Ethics approval

All animal experiments were conducted in accordance with institutional as well as national guidelines and regulations and were approved by the Stanford University Administrative Panel on Laboratory Animal Care (APLAC), an AAALAC-accredited program (protocol no. 34459).

### Animals and housing

C57BL/6 mice were used in all experiments. Male mice were used for all primary cohorts. Female mice were included only in the young LPS model, and these data are reported in the Supplementary Information. Young mice were 2–3 months old, aged mice were 18–20 months old, and super-aged mice were 26–28 months old at the start of experiments. Young and aged mice were purchased from The Jackson Laboratory, and super-aged mice were obtained from the National Institute on Aging (NIA) Aged Rodent Colony and acclimated in-house under identical husbandry conditions for 1–2 weeks prior to experiments.

Animals were housed under identical husbandry conditions with ad libitum access to food and water, maintained on a 12-h light/dark cycle (lights on at 07:00 and off at 19:00) with controlled temperature of 20-26°C and humidity of 30-60%. Before surgery, mice were group-housed (4–5 mice per Innocage®). After EEG head-mount implantation, mice were singly housed to prevent damage to implants.

Sample sizes were estimated based on effect sizes observed in our prior studies using the same EEG and behavioral platforms [25, 26]. Planned and final sample sizes for the BSEEG and survival cohorts are provided in Supplementary Table 1. Animals were allocated to vehicle or zuranolone-treated groups using weight-stratified randomization.

### Zuranolone formulation and administration

Zuranolone (>98% by HPLC; AOBIOUS, Inc.) was supplied as a powder and dissolved in 30% (w/v) sulfobutylether-β-cyclodextrin (SBE-β-CD; 98%; MedChemExpress) immediately before use. Zuranolone or vehicle (30% SBE-β-CD) was administered by oral gavage using a sterile, disposable, flexible animal feeding needle (20G, 1.5 in.; Fisher Scientific). Across experiments, zuranolone was administered at 0.5 or 1.0 mg kg^−1^ depending on the model, age group, and dosing paradigm. Detailed dosing schedules for the primary BSEEG and survival cohorts, including dose, timing, and prevention or treatment paradigm, are provided in Supplementary Table 1; additional supplemental cohorts are described in the corresponding supplementary figure legends.

### EEG electrode head-mount placement surgery

EEG electrode head-mount placement surgery was performed as described previously [13]. Mice were anesthetized with isoflurane (1–3% inhalation), and the skull was exposed. A stainless-steel EEG head mount (#8201; Pinnacle Technology, Inc.) was positioned with its center aligned to the midline of the skull, with anterior holes located 2 mm anterior to bregma and posterior holes 2 mm anterior to lambda (±2 mm mediolateral). Holes were drilled using a 23-gauge needle, and stainless-steel screws (anterior: #8209; posterior: #8212) were inserted. The head mount was secured with dental cement. Postoperative analgesia was provided via subcutaneous meloxicam (5 mg kg^−1^).

### EEG recording and BSEEG signal processing

Following recovery, mice were tethered to a continuous EEG recording system (#8200-K1-SL; Pinnacle Technology, Inc.). EEG signals were recorded from the right frontal cortex (EEG2) and right parietal cortex (EEG1), with the left frontal screw serving as ground and the left parietal screw as reference. Based on prior validation [25, 26], EEG2 was used for all analyses.

Signals were acquired using Sirenia® software and exported as EDF files. Power spectral density was calculated using fast Fourier transformation, and BSEEG scores were derived as the ratio of 3-Hz to 10-Hz power, consistent with our previously described and validated method [27] [28]. Thus, a higher BSEEG score represents a shift toward low-frequency power, consistent with delirium in humans and a delirium-like condition in mice. Raw EEG data were processed using an updated version of our publicly available web-based analysis tool (https://sleephr.jp/gen/mice7days/) [29], which calculated hourly averaged BSEEG scores. For visualization and statistical analyses, hourly BSEEG values were further averaged into 12-h bins corresponding to daytime and nighttime periods. Group-averaged standardized BSEEG (sBSEEG) time courses are shown in Fig. 1, and individual-mouse sBSEEG time courses are provided in Supplementary Fig. 1–11.

**Figure 1.**
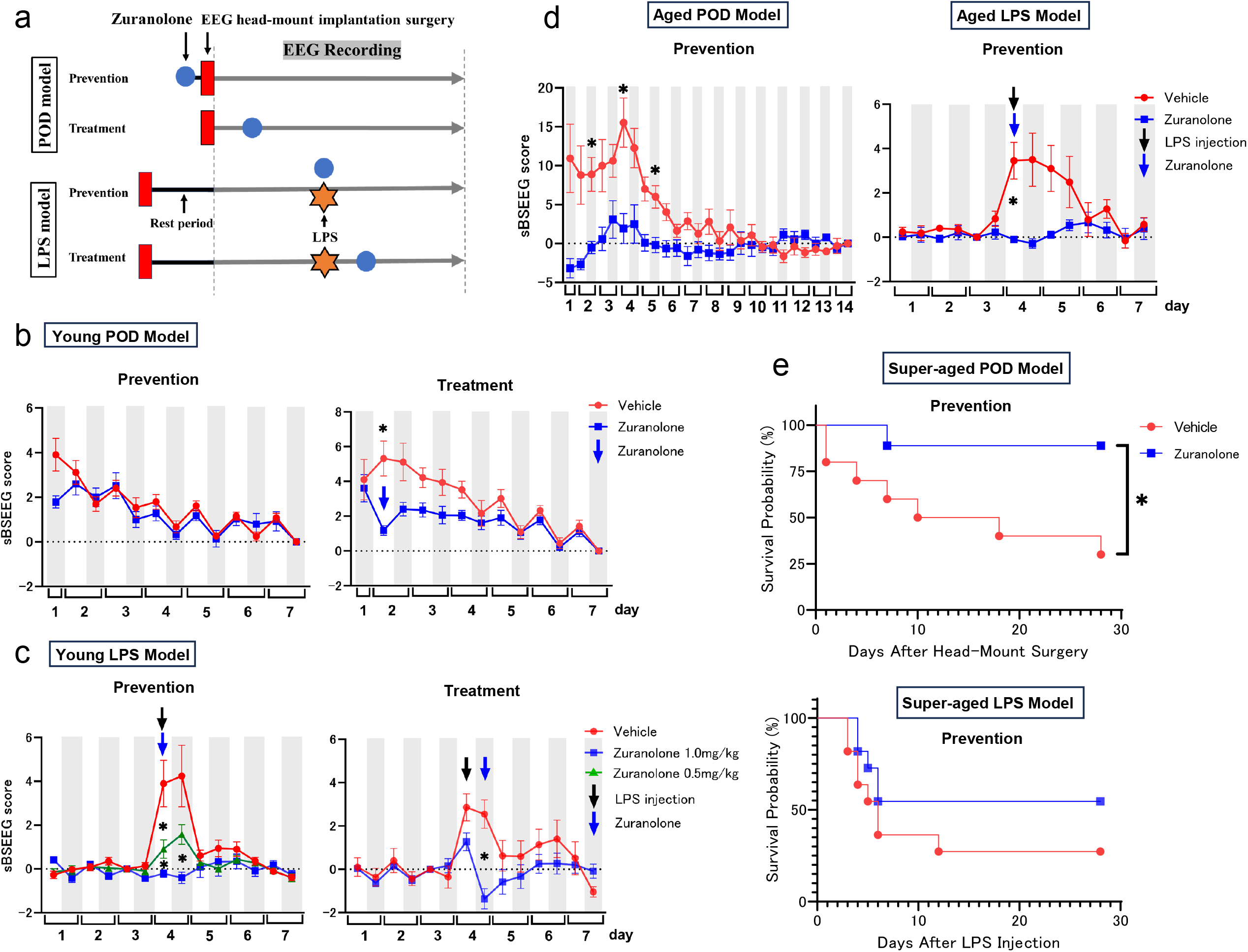
Zuranolone attenuates POD- and LPS-associated increases in standardized BSEEG (sBSEEG) scores across age cohorts. **(a)** Experimental timelines. Mice underwent either the postoperative delirium (POD) model induced by EEG head-mount implantation surgery or the LPS-induced systemic inflammation model. Zuranolone or vehicle was administered using prevention or treatment paradigms, followed by continuous EEG recording. For the LPS model, administration immediately before LPS injection was defined as a prevention/early-intervention paradigm. **(b)** Young POD model. Time course of sBSEEG scores following surgery under prevention (pre-surgery) and treatment (post-surgery) paradigms. **(c)** Young LPS model. Time course of sBSEEG scores following LPS challenge, demonstrating dose-dependent suppression of LPS-associated EEG abnormalities by zuranolone under prevention and treatment paradigms. **(d)** Aged cohort. Time courses of sBSEEG scores in the POD and LPS models under prevention dosing. **(e)** Super-aged cohort. Kaplan–Meier survival curves following POD induction or LPS challenge under prevention dosing. For panels (b–d), sBSEEG scores were calculated from EEG data acquired within predefined analysis windows and are shown as mean ± SEM. Grey shading indicates nighttime periods. Asterisks denote time-bin–specific differences between vehicle and zuranolone within the predefined analysis window, tested using a mixed-effects model with Šidák correction for multiple comparisons (adjusted P < 0.05). For panel (e), survival curves were analyzed using Kaplan–Meier analysis and compared using the log-rank test. Statistical models, handling of repeated measures, and multiple-comparison corrections are described in Methods. Group sizes, dosing, and administration schedules are provided in Methods and Supplementary Table 1. Individual-mouse sBSEEG time courses for the young and aged cohorts are shown in Supplementary Fig. 1–4 and 8–9, and those for the super-aged cohorts are shown in Supplementary Fig. 10–11.

### EEG recording duration and predefined analysis windows

Continuous EEG recordings were performed for up to 7 days in all experimental cohorts. In the aged POD cohort, EEG recordings were extended to 14 days to characterize prolonged recovery phase associated with advanced age. In the POD model, the primary analysis window was defined as 0–96 hours (four days) after surgery, corresponding to the acute postoperative phase during which delirium-like EEG abnormalities (BSEEG increase) are known to peak [30]. In the LPS model, the primary analysis window was defined as 0–48 hours (2 days) after LPS administration, encompassing the period of maximal inflammation-associated EEG slowing (increased BSEEG score). EEG data acquired outside these primary analysis windows were used descriptively to visualize longer-term recovery trajectories and confirm return toward baseline, and were not included in the primary statistical analyses.

### Standardized BSEEG (sBSEEG) score

To facilitate within-animal comparisons and highlight deviations from a reference state, we calculated standardized BSEEG (sBSEEG) scores as 12-h mean BSEEG values referenced to a prespecified baseline for each mouse, consistent with our prior work [25, 26]. Daytime and nighttime periods were defined by the 07:00/19:00 light/dark cycle.

In the POD model, baseline EEG cannot be obtained prior to the EEG head-mount implantation surgery. Therefore, for each mouse, the baseline was defined as the nighttime 12-h mean BSEEG score on the final recording day, which was assumed to approximate a postoperative steady state within the recording duration; this reference bin was defined as sBSEEG = 0. This approach is consistent with prior validation demonstrating stabilization of BSEEG values by this time point following head-mount implantation surgery [26].

In the LPS model, baseline EEG was available before the inflammatory trigger. Thus, for each mouse, the baseline was defined as the daytime 12-h mean BSEEG score on the day preceding LPS injection, which was defined as sBSEEG = 0, and subsequent bins reflect deviations from this pre-insult baseline.

### Experimental models

Detailed dosing schedules for the primary prevention and treatment paradigms are provided in Supplementary Table 1, with additional supplemental cohorts described in the corresponding supplementary figure legends.

#### LPS model

Systemic inflammation was induced by intraperitoneal injection of lipopolysaccharide (LPS; *Escherichia coli* O111:B4, Sigma-Aldrich). Young mice received 1.0 mg kg^−1^, aged mice 0.1 mg kg^−1^, and super-aged mice 0.05 mg kg^−1^ to account for increased age-dependent susceptibility [31]. Mice were allowed to recover for at least 2 weeks after EEG head-mount implantation before EEG recording. Zuranolone or vehicle was administered according to prevention or treatment paradigms. In the prevention/early-intervention paradigm, zuranolone or vehicle was administered immediately before LPS injection; in the treatment paradigm, dosing was initiated 6 hours after LPS injection.

#### POD model

Surgical stress associated with EEG head-mount implantation served as the delirium-inducing insult. For prevention dosing, zuranolone or vehicle was administered by oral gavage 10 min before surgery; for treatment dosing, zuranolone or vehicle was administered after surgery. EEG recording commenced after surgery and continued continuously for the duration of the experiment.

### Behavioral assessments

Behavioral testing was performed only in young mice during the light phase using a standardized battery assessing attention, arousal, and working memory, including the buried food test, open-field test, and Y-maze test. Behavioral testing was conducted at baseline and at 24 h after surgery or LPS administration. Behavioral performance was quantified using video tracking and analyzed using SMART 3.0 software.

Composite Z scores were calculated from the behavioral test battery as a quantitative index of delirium-like behavioral severity in mice, aligned with core domains assessed by the Confusion Assessment Method (CAM), as previously described [25, 26]. The behavioral battery comprised six parameters: (i) latency to eat food, (ii) time spent in the center, (iii) latency to enter the center, (iv) freezing time, (v) entries into the novel arm, and (vi) duration in the novel arm. For each behavioral parameter, the change from baseline (ΔX) was calculated for each mouse. Individual Z scores were computed as Z = [ΔX_test − mean(ΔX_control)] / SD(ΔX_control), where ΔX_control represents the baseline-corrected behavioral change in the corresponding young vehicle-treated control group.

The composite Z score for each mouse was calculated as the sum of the six individual Z scores and normalized by the standard deviation of the summed Z scores in the corresponding control group. Before summation, individual behavioral Z scores were directionally adjusted as needed so that higher values consistently reflected greater behavioral impairment. Because behavioral experiments were conducted exclusively in young mice, reference distributions were derived from baseline-corrected behavioral changes in young vehicle-treated control groups within each experimental model, defined as surgery + vehicle for the POD model and LPS + vehicle for the LPS model. Model-specific reference definitions are detailed in the Supplementary Methods. Individual Z scores for each behavioral parameter and composite Z scores for all animals are provided in Supplementary Tables 2 (POD) and 3 (LPS).

### Immunofluorescence staining and quantification

To assess the cellular evidence of neuroinflammation, mice were deeply anesthetized with isoflurane and transcardially perfused with saline followed by 4% periodate–lysine–paraformaldehyde fixative. Brains were extracted and coronally sectioned at 40 μm thickness to include hippocampal and cortical regions.

Free-floating sections were blocked with 5% bovine serum albumin for 1 h and incubated overnight at 4 °C with primary antibodies against IBA1 (1:400; FUJIFILM Wako) and CD68 (1:400; Cell Signaling Technology). After washing, sections were incubated with fluorophore-conjugated secondary antibodies (Alexa Fluor® 488 or 594; 1:500; Thermo Fisher Scientific) for 2 h at room temperature. Nuclei were counterstained with DAPI (Vector Laboratories), and sections were mounted with Vectashield mounting medium.

Images were acquired using a fluorescence microscope (BZ-X800, KEYENCE) under identical acquisition settings across groups. Quantitative analyses of microglial density and activation were performed using BZ-X800 Analyzer software across 8–10 sections per mouse.

### Statistical analysis

Statistical analyses were performed using GraphPad Prism 9. Data normality was assessed using the Shapiro–Wilk test. Two-group comparisons were conducted using Welch’s t-test or the Mann– Whitney U test, as appropriate. For multi-group comparisons, Kruskal–Wallis tests were performed followed by Dunn’s multiple-comparisons tests for prespecified pairwise contrasts. Dunn-adjusted P values were used to determine significance for these post hoc comparisons.

For longitudinal BSEEG analyses, we used a mixed-effects model for repeated measures within the predefined analysis windows (POD: 0–96 h post-surgery; LPS: 0–48 h post-injection). Planned time-bin–specific comparisons between vehicle and zuranolone were performed from the fitted model, with Šidák correction for multiple comparisons across time bins within each model/cohort and planned comparison; Šidák-adjusted P values were used to determine significance for these time-bin–specific comparisons. Survival curves were analyzed using Kaplan–Meier analysis and compared using the log-rank test. Data are presented as mean ± SEM. Nominal two-sided P < 0.05 was considered statistically significant for tests without multiple-comparison adjustment, whereas adjusted P < 0.05 was considered significant for Dunn’s and Šidák-corrected comparisons.

## Results

### Zuranolone attenuates sBSEEG increases in the POD and LPS models across age cohorts (Fig. 1)

To determine whether zuranolone ameliorates delirium-like electrophysiological dysfunction, we quantified changes in BSEEG scores in the POD and LPS models across young, aged, and super-aged cohorts (Fig. 1a; dosing and schedules in Supplementary Table 1). Group-averaged sBSEEG trajectories are shown in Fig. 1, and corresponding individual-mouse trajectories are provided in Supplementary Fig. 1–11. Neither the vehicle (SBE-β-CD) nor zuranolone alone produced appreciable changes in baseline sBSEEG scores (Supplementary Fig. 7).

In the young POD model, postoperative elevations in BSEEG scores were observed during the acute phase following surgery (Fig. 1b). Prophylactic (preoperative) administration of zuranolone did not significantly reduce BSEEG scores across the overall 0–96 h post-surgery analysis window compared with vehicle. However, during the first postoperative night, BSEEG scores were lower in the prophylactic zuranolone group than in the vehicle group, although this difference did not reach statistical significance (P = 0.13). In contrast, zuranolone administered after surgery in a treatment paradigm significantly reduced postoperative BSEEG elevations relative to vehicle, indicating efficacy when given after the onset of POD-associated electrophysiological abnormalities.

Systemic LPS administration in young mice induced marked increases in BSEEG scores and disruption of diurnal EEG patterns (Fig. 1c). Preventive administration of zuranolone significantly attenuated LPS-induced BSEEG elevations in a dose-dependent manner, with greater suppression observed at the higher dose (1.0 mg/kg) compared with the lower dose (0.5 mg/kg). Comparable preventive effects were observed in young female mice (Supplementary Fig. 6). Similarly, treatment administration of zuranolone after LPS injection significantly reduced BSEEG elevations compared with vehicle, supporting its efficacy during the acute inflammatory phase. In an additional cohort, we tested a combined dosing schedule in which the same mice received zuranolone immediately before LPS injection (prevention/early-intervention) and again 24 h after LPS injection (treatment), using a lower dose (0.5 mg/kg) given the two consecutive administrations, in contrast to the single dose prevention- or treatment-only paradigms (1.0 mg/kg). The experimental timeline, group-averaged sBSEEG trajectory, and individual-mouse traces are shown in Supplementary Fig. 5. In this combined dosing cohort, a similar suppression of LPS-associated sBSEEG elevations was observed, with a shorter duration of elevation compared with vehicle.

In the aged cohorts, both the POD and LPS models produced larger and more prolonged BSEEG increases than those observed in young mice (Fig. 1d). Under the prevention paradigm, zuranolone significantly reduced BSEEG elevations in both the aged POD and aged LPS models, indicating that zuranolone retained efficacy in ameliorating the electrophysiological disturbances despite this age-related vulnerability.

Finally, in the super-aged cohort, POD induction by head-mount implantation surgery was associated with substantial mortality (Fig. 1e). Prophylactic zuranolone significantly improved survival relative to vehicle. In the super-aged LPS model, although the between-group difference in survival did not reach statistical significance (P = 0.18), survival was directionally higher in the zuranolone group. The corresponding sBSEEG trajectories for the super-aged POD and LPS prevention cohort are provided in Supplementary Fig. 10 and 11.

Together, these results indicate that zuranolone attenuated delirium-like BSEEG abnormalities across POD and LPS models, with efficacy varying by age group, model, and dosing timing. Zuranolone also improved survival after POD induction in super-aged mice, whereas prophylactic efficacy in the young POD model was limited to an early postoperative trend.

### Zuranolone improves delirium-like behavioral outcomes in young mice, with stronger effects in the LPS model (Fig. 2)

We next tested whether electrophysiological improvement was accompanied by mitigation of delirium-like behavioral deficits using a standardized battery assessing motivated behavior/attention (buried food test), arousal and exploration (open-field test), and working memory (Y-maze).

**Figure 2.**
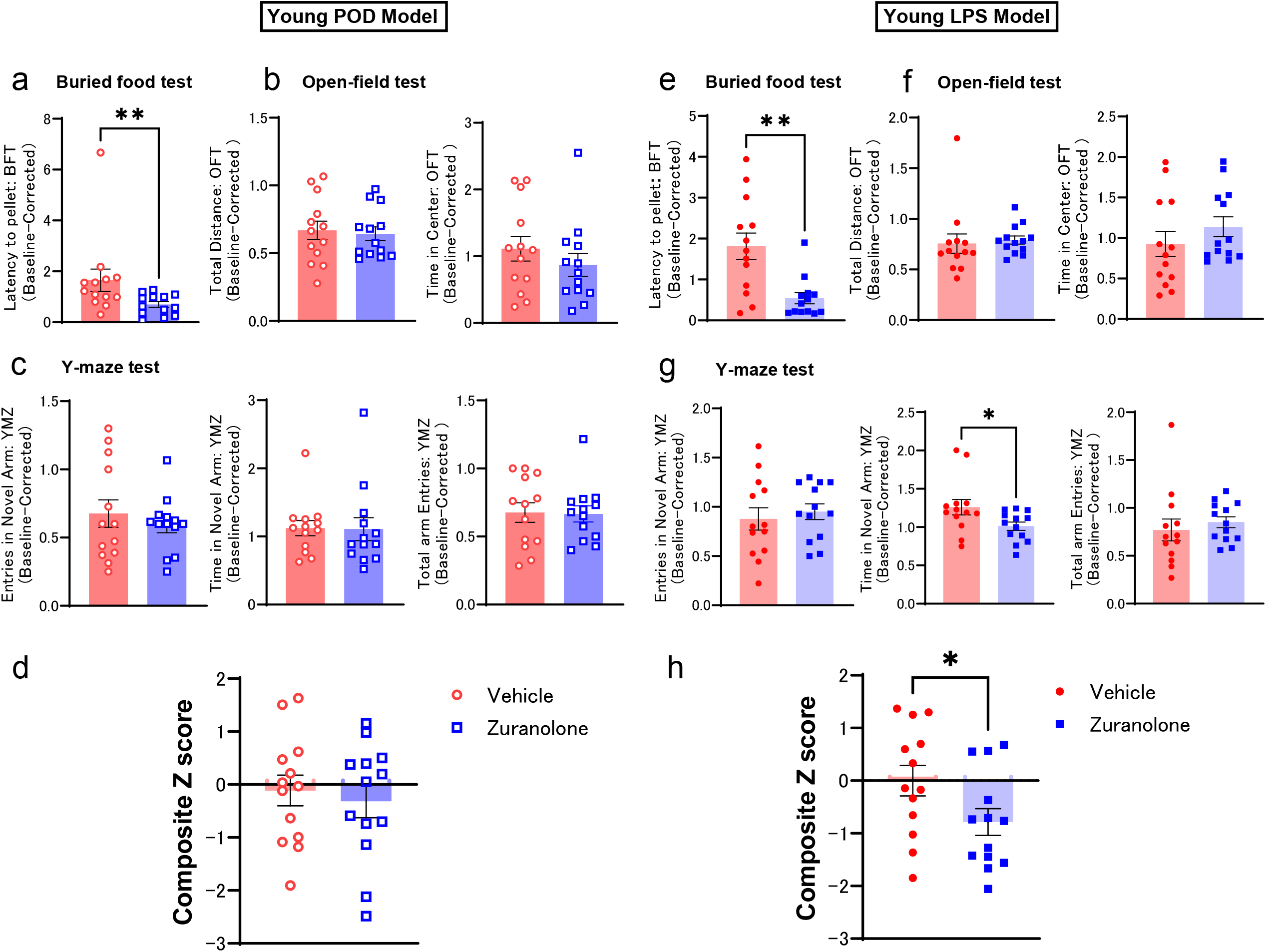
Zuranolone improves delirium-like behavioral outcomes in young mice, with stronger effects in the LPS model. Behavioral outcomes were assessed at baseline and 24 h after the insult and plotted as baseline-corrected values (post − pre). Behavioral variables were sign-adjusted for composite Z score calculation so that higher values consistently reflect greater impairment (see Methods). Each point represents one mouse; bars indicate mean ± SEM. (a–d) Young POD model. (a) Buried food test (latency). (b) Open-field test (total distance traveled and time spent in center). (c) Y-maze test (novel-arm entries and time spent in the novel arm, and total arm entries). (d) Composite Z score integrating the behavioral battery (higher values indicate greater impairment). (e–h) Young LPS model. (e) Buried food test. (f) Open-field test. (g) Y-maze test. (h) Composite Z score. Statistical annotations: *P < 0.05, **P < 0.01 (tests as described in Methods).

Outcomes were baseline-corrected (post − pre) and integrated into a composite behavioral Z score (Methods).

In the young POD model, zuranolone significantly improved performance in the buried food test, reflected by reduced baseline-corrected latency to retrieve the pellet compared with vehicle (Fig. 2a). In contrast, open-field outcomes (baseline-corrected total distance traveled and time spent in the center) did not significantly differ between groups (Fig. 2b). Y-maze measures of novel-arm exploration and total entries were also not significantly altered (Fig. 2c). Consistent with these mixed domain-specific effects at 24 h, the composite behavioral Z score did not significantly differ between vehicle- and zuranolone-treated mice in the POD model (Fig. 2d).

In the young LPS model, zuranolone significantly improved buried food test performance (Fig. 2e). Open-field measures did not show significant differences (Fig. 2f). In the Y-maze, zuranolone-treated mice exhibited a significant difference in time spent in the novel arm relative to vehicle-treated mice, while other Y-maze measures—including novel-arm entries and total arm entries—were not significantly changed (Fig. 2g).

When behavioral outcomes across domains were integrated using the composite Z score, zuranolone significantly reduced overall behavioral impairment following LPS challenge (Fig. 2h), indicating a global attenuation of delirium-like behavioral severity in the inflammation-driven model despite heterogeneous effects at the level of individual behavioral metrics.

Together, these behavioral data indicate that zuranolone mitigated delirium-like behavioral impairment in young mice following systemic inflammatory challenge, with the most robust treatment effect captured by the composite behavioral severity metric. In contrast, behavioral effects in the young POD model were limited and domain-specific at the 24-h time point.

### Zuranolone reduces surgery-associated microglial activation and attenuates LPS-induced microglial responses in young mice (Fig. 3)

We next quantified IBA1+ and CD68+ microglial measures by immunofluorescence in the POD and LPS models. Brains were collected 24 h after head-mount implantation surgery in the POD model and 24 h after LPS injection in the LPS model for immunostaining (Methods).

**Figure 3.**
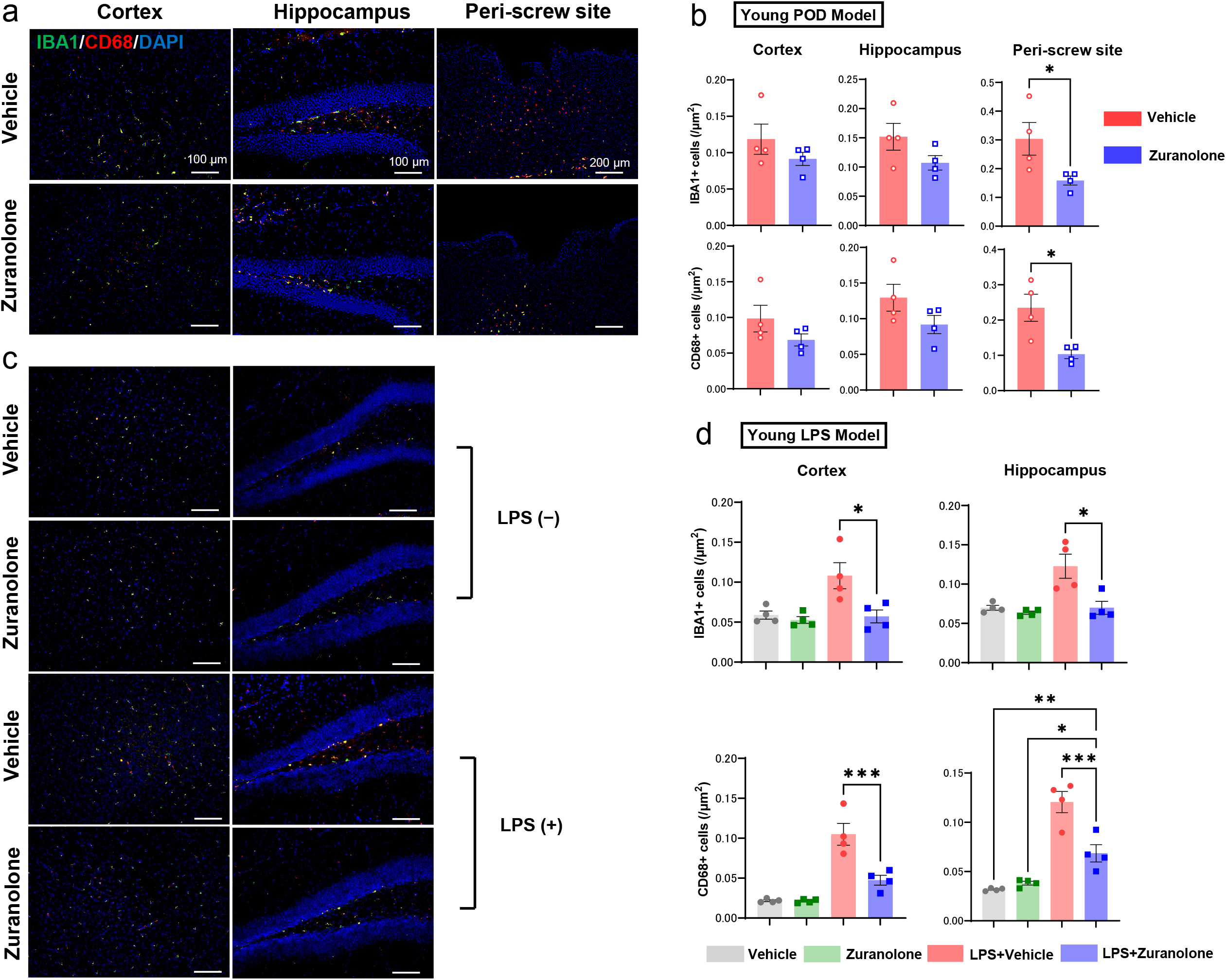
Zuranolone reduces surgery-associated microglial activation and attenuates LPS-induced microglial responses in young mice. Immunofluorescence staining for IBA1 (microglia; green) and CD68 (activation marker; red) with DAPI (nuclei; blue). Each point represents one mouse; bars indicate mean ± SEM. (a) Representative images from cortex, hippocampus, and peri–screw site region in the young POD model (vehicle vs zuranolone). Scale bars, 100 μm (cortex and hippocampus) and 200 μm (peri–screw site). (b) Quantification of IBA1+ and CD68+ signal in cortex, hippocampus, and peri–screw site in the young POD model. (c) Representative cortical and hippocampal images in the young LPS model under LPS(−) and LPS(+) conditions with vehicle or zuranolone. Scale bars as indicated. (d) Quantification of IBA1+ and CD68+ signal in cortex and hippocampus in the young LPS model across groups (vehicle, zuranolone, LPS + vehicle, LPS + zuranolone), demonstrating LPS-driven increases and attenuation by zuranolone. Statistical annotations: *P < 0.05, **P < 0.01, ***P < 0.001. Quantification and statistical methods, including Dunn’s multiple comparisons for prespecified pairwise contrasts, are described in Methods.

In the young POD model, vehicle-treated mice showed prominent IBA1 and CD68 signal, particularly in the peri–screw site region adjacent to the implantation site (Fig. 3a). Quantification revealed that zuranolone significantly reduced both IBA1+ and CD68+ measures at the peri-screw site compared with vehicle (Fig. 3b). In the hippocampus, reductions showed a trend for IBA1 (P = 0.11) and were not significant for CD68, whereas cortical measures did not significantly differ between groups. These results indicate that zuranolone suppresses surgery-associated microglial activation particularly at the peri-screw site in young mice.

In the young LPS model, LPS administration increased microglial measures in both cortex and hippocampus, reflected by elevated IBA1+ density and CD68+ signal under LPS(+) conditions (Fig. 3c,d). Zuranolone had minimal effects under LPS(−) conditions. Under LPS(+) conditions, zuranolone significantly attenuated LPS-induced increases in IBA1 and CD68 in both cortex and hippocampus compared with LPS + vehicle (Fig. 3d; Kruskal–Wallis followed by Dunn’s multiple comparisons for prespecified pairwise contrasts).

Together, these findings show that zuranolone reduced microglial activation 24 h after POD surgery, particularly at the peri-screw site, and dampened inflammation-driven microglial responses 24 h after LPS.

### Zuranolone suppresses microglial activation in the aged POD and LPS models (Fig. 4)

To test whether these neuroimmune effects extend to older animals, we performed the same immunofluorescence analyses in aged mice, collecting brains 24 h after surgery (aged POD) or 24 h after LPS injection (aged LPS).

**Figure 4.**
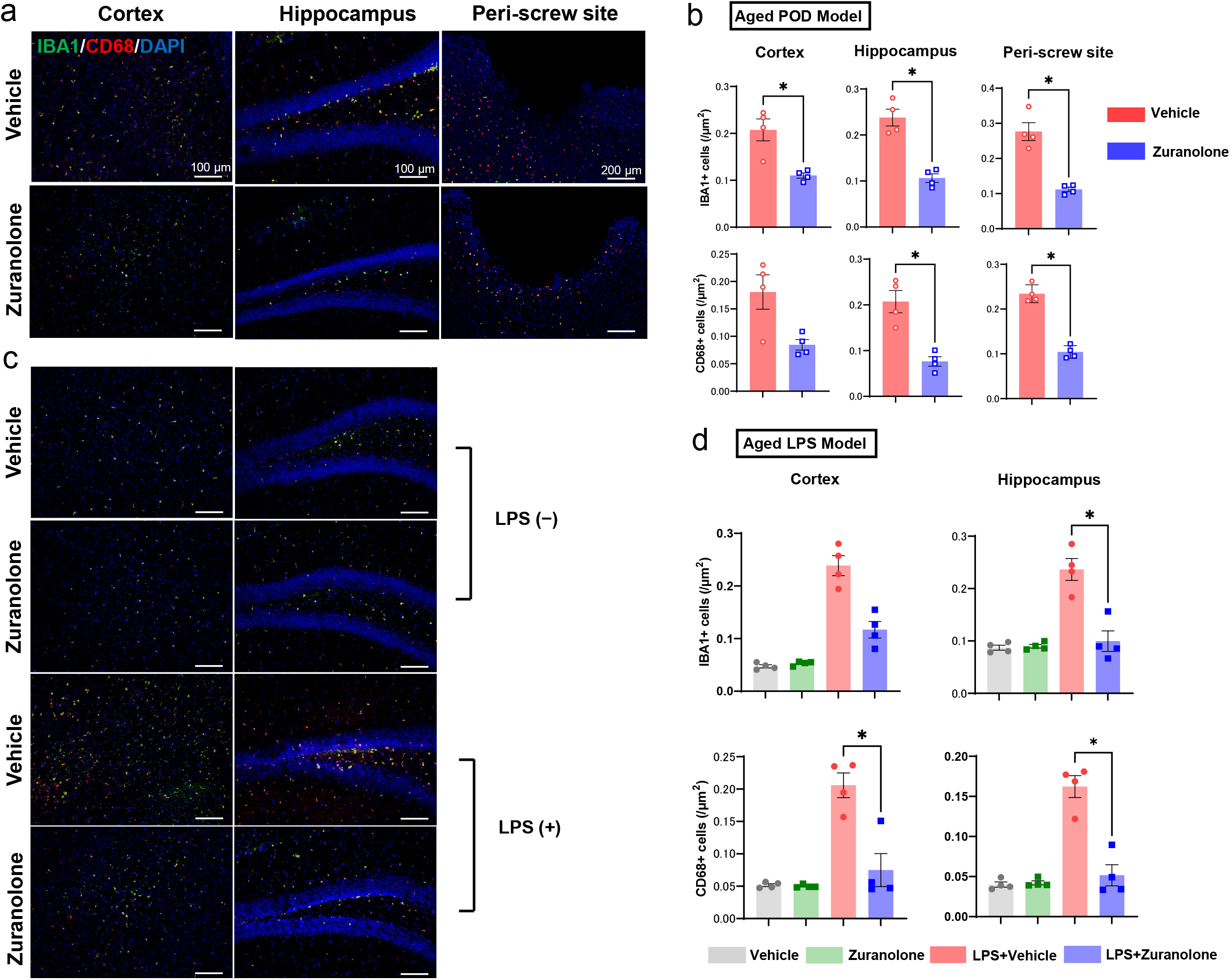
Zuranolone suppresses POD- and LPS-associated microglial activation in aged mice. Immunofluorescence staining for IBA1 (microglia; green) and CD68 (activation marker; red) with DAPI (nuclei; blue). Each point represents one mouse; bars indicate mean ± SEM. (a) Representative images from cortex, hippocampus, and peri–screw site region in the aged POD model (vehicle vs zuranolone). Scale bars, 100 μm (cortex and hippocampus) and 200 μm (peri– screw site). (b) Quantification of IBA1+ and CD68+ signal in cortex, hippocampus, and peri–screw site in the aged POD model. (c) Representative cortical and hippocampal images in the aged LPS model under LPS(−) and LPS(+) conditions with vehicle or zuranolone. Scale bars as indicated. (d) Quantification of IBA1+ and CD68+ signal in cortex and hippocampus in the aged LPS model across groups (vehicle, zuranolone, LPS + vehicle, LPS + zuranolone). Statistical annotations: *P < 0.05. Quantification was performed under identical acquisition settings across groups; statistical methods, including Dunn’s multiple comparisons for prespecified pairwise contrasts, are described in Methods.

In the aged POD model, vehicle-treated mice displayed marked microglial signal across regions, with prominent activation at the peri–screw site (Fig. 4a). Quantitatively, zuranolone significantly reduced IBA1+ measures in cortex, hippocampus, and peri-screw site (Fig. 4b). For CD68, zuranolone significantly reduced signal in the hippocampus and peri-screw site, while cortical CD68 did not differ significantly (P = 0.11). These data indicate that in aged mice, zuranolone broadly suppresses surgery-associated microglial activation across multiple brain regions.

In the aged LPS model, LPS administration increased microglial density/activation measures across regions (Fig. 4c,d). Zuranolone did not significantly alter measures under LPS(−) conditions. Under LPS(+) conditions, zuranolone significantly attenuated LPS-induced microglial responses, including reductions in CD68 signal in both cortex and hippocampus and IBA1+ density in the hippocampus, with consistent directional attenuation across all examined measures (Fig. 4d; Kruskal–Wallis followed by Dunn’s multiple comparisons for prespecified pairwise contrasts).

Collectively, these data indicate that zuranolone robustly attenuated 24-h post-insult microglial activation in aged mice after both POD surgery and LPS challenge, with broadly consistent effects across the examined brain regions.

## Discussion

We demonstrate that the neuroactive steroid zuranolone mitigates delirium-like brain dysfunction across electrophysiological, neuroimmune, and behavioral domains. We observe these effects across young, aged, and super-aged mice in models of postoperative stress and systemic inflammation.

Continuous EEG quantified by the BSEEG method provided a chronologically precise readout of delirium-like brain dysfunction, while behavioral testing and regional microglial analyses provided phenotypic and mechanistic context that aligned with the EEG findings. Across both the POD and LPS models, the magnitude and persistence of delirium-like abnormalities increased with age, and zuranolone’s efficacy depended on the type of insult (surgical vs inflammatory) and the timing of administration.

### Model-dependent efficacy suggests a strong impact on inflammation-driven brain dysfunction

A key finding was the robust suppression of LPS-associated BSEEG increases, reflecting low-frequency EEG dominance, by zuranolone, including a dose-dependent effect observed in young mice and similarly preserved efficacy in aged mice. A similar effect was also evident in young female mice. In parallel, zuranolone improved delirium-like behavioral performance after LPS challenge, with the strongest global effect captured by the composite behavioral severity metric. The LPS paradigm elicits a rapidly evoked systemic inflammatory response that drives acute brain dysfunction, reflected by pronounced BSEEG elevations [8] [9] [25].

In contrast to the inflammation-driven LPS paradigm, the postoperative model highlighted a stronger dependence on dosing time and baseline vulnerability. In the young POD model, a single prophylactic dose did not significantly reduce postoperative BSEEG abnormalities across the acute window (0–96 h), whereas post-surgery treatment significantly attenuated postoperative BSEEG elevations. This discrepancy is consistent with relatively low baseline susceptibility in young animals—potentially reflecting limited age-related microglial priming and preserved endogenous neurosteroid tone—such that benefit is more apparent when dosing is aligned with the emergence of postoperative EEG abnormalities rather than delivered as a single preoperative intervention. By contrast, in older animals, heightened baseline neuroimmune and electrophysiological reactivity may increase vulnerability to surgical stress and make prophylactic benefit more readily detectable. In parallel, aging has been associated with reduced brain allopregnanolone levels and lower 5α-reductase activity, supporting the possibility that altered endogenous neurosteroid signaling may contribute to age-dependent susceptibility to delirium-like brain-state disturbances [32]. More broadly, these findings support a brain-state and timing-dependent model of pharmacological benefit, with translational implications for aligning drug exposure with predictable high-risk periods rather than assuming earlier administration is uniformly superior.

Together, these findings indicate that delirium-like abnormalities vary by insult type, age, and timing of intervention, and that modulation of neurosteroid signaling may be particularly effective when systemic inflammation or age-related vulnerability is a primary driver.

### Aging increases susceptibility and reveals prophylactic benefit

Across paradigms, aged mice showed larger and more prolonged BSEEG abnormalities than young mice, consistent with reduced physiological reserve and heightened baseline inflammatory tone in aging [6] [13] [25]. Under these higher-vulnerability conditions, prophylactic zuranolone reduced BSEEG elevations in both aged POD and LPS models, indicating maintained efficacy in ameliorating delirium-associated electrophysiological disturbances and suggesting that prophylactic benefit becomes more apparent as vulnerability increases with age.

Extending this age gradient to the super-aged cohort, prophylactic zuranolone significantly improved survival after surgical insult. In the super-aged LPS model, survival after LPS challenge was not significantly improved, but the direction of effect was consistent with the significant survival benefit observed after surgical insult.

Together, these findings mirror clinical observations that delirium risk and adverse outcomes increase with age and support the view that preventive strategies may yield the largest benefit in high-risk populations [6] [7].

### Neuroimmune effects provide mechanistic context

Microglial analyses at 24 h after insult provided mechanistic context for zuranolone’s electrophysiological and behavioral effects. In young POD mice, zuranolone most strongly attenuated microglial activation at the peri–screw site, consistent with modulation of local injury-associated neuroinflammation. In aged POD mice, suppression extended across cortex, hippocampus, and the peri–screw site, supporting the concept that identical surgical stress elicits a more diffuse and amplified neuroimmune response with age [8] [15]. In the LPS model, zuranolone had minimal impact under non-inflammatory conditions but significantly reduced LPS-induced microglial activation in cortex and hippocampus in young mice, and also significantly attenuated LPS-induced microglial responses in aged mice, with effects spanning cortical and hippocampal regions, consistent with context-dependent dampening of inflammation-evoked neuroimmune responses. Our findings align with prior work in these same paradigms showing that BSEEG correlates with delirium-like behavioral deficits and with microglial activation after LPS challenge. These studies further suggest that BSEEG provides a sensitive electrophysiological readout of postoperative brain dysfunction, particularly in settings where behavioral assessments become less reliable due to aging or increased surgical burden [25] [26]. Taken together, these results support an association between neuroimmune activation and delirium-like EEG abnormalities quantified by BSEEG, and motivate future studies that directly test causal mechanisms (e.g., time-resolved microglial profiling, or microglia-targeted interventions).

### Electrode-adjacent (peri–screw site) microglial responses as a focal readout of implant-associated neuroinflammation

Intracranial electrode implantation is known to elicit a stereotyped foreign-body response in which microglia in proximity to the implant become activated shortly after surgery, followed over subsequent hours to days by recruitment/migration of additional microglia and release of pro-inflammatory mediators [33] [34] [35]. In this context, the peri–screw site region provides a sensitive, anatomically anchored compartment to quantify implant-associated neuroimmune activation and its modulation by therapy. Our finding that zuranolone reduced IBA1/CD68 immunoreactivity at the peri–screw site at 24 h after POD surgery is therefore consistent with attenuation of the early implant-associated microglial response that may otherwise propagate into broader recruitment and inflammatory amplification over time.

### Limitations and future directions

Several limitations warrant consideration. First, behavioral testing was performed at discrete time points and therefore cannot fully capture the hallmark fluctuation of delirium-like states [20]. Second, microglial analyses were conducted at a single post-insult time point, limiting inference about the temporal dynamics of neuroinflammation and its relationship to evolving EEG abnormalities. In particular, with a single 24-h endpoint, we cannot distinguish effects on immediate peri–screw site microglial activation from effects on subsequent recruitment or migration dynamics described in electrode implantation paradigms [33] [34] [35]. Third, we did not directly measure plasma or brain concentrations of zuranolone in these paradigms, and dose–response relationships were not systematically evaluated across all models, age groups, and treatment schedules. These limitations restrict PK/PD inference regarding the optimal dose, timing, and duration of exposure across age groups and insult types. Fourth, female mice were included only in the young LPS model, where BSEEG analysis similarly demonstrated significant suppression of LPS-associated abnormalities by zuranolone (Supplementary Fig. 6). Nevertheless, sex-specific effects on behavioral or neuroimmune outcomes, as well as effects in aged or postoperative models, remain to be determined. Finally, the POD model uses head-mount implantation surgery as the inducing insult; while it reliably produces delirium-like EEG and behavioral changes in prior work, it represents a specific surgical and anesthetic context, and future studies should test generalizability across procedures and perioperative conditions [12] [13] [25] [26].

Despite these limitations, our findings demonstrate that zuranolone mitigates delirium-like electrophysiological abnormalities across surgical and inflammatory insults and across age cohorts, with particularly robust effects in inflammation-driven and age-vulnerable contexts. These data support a framework in which enhancing inhibitory neurosteroid signaling can stabilize acute pathological brain states relevant to delirium and motivate further work to optimize dosing strategies, identify responsive endotypes, and test translational potential in clinically aligned settings to ultimately improve patient outcomes.

## Data Availability

The datasets generated and/or analyzed in this study are available from the corresponding author upon reasonable request.

## References

1. Goldberg, T.E., et al., Association of Delirium With Long-term Cognitive Decline: A Meta-analysis. JAMA Neurol, 2020. 77(11): p. 1373–1381.

2. Leslie, D.L., et al., One-year health care costs associated with delirium in the elderly population. Arch Intern Med, 2008. 168(1): p. 27–32.

3. Thom, R.P., et al., Delirium. Am J Psychiatry, 2019. 176(10): p. 785–793.

4. Prendergast, N.T., P.J. Tiberio, and T.D. Girard, Treatment of Delirium During Critical Illness. Annu Rev Med, 2022. 73: p. 407–421.

5. Bramley, P., et al., Risk factors for postoperative delirium: An umbrella review of systematic reviews. Int J Surg, 2021. 93: p. 106063.

6. Inouye, S.K., R.G. Westendorp, and J.S. Saczynski, Delirium in elderly people. Lancet, 2014. 383(9920): p. 911–22.

7. Fong, T.G. and S.K. Inouye, The inter-relationship between delirium and dementia: the importance of delirium prevention. Nat Rev Neurol, 2022. 18(10): p. 579–596.

8. Hoogland, I.C., et al., Systemic inflammation and microglial activation: systematic review of animal experiments. J Neuroinflammation, 2015. 12: p. 114.

9. Kimchi, E.Y., et al., Clinical EEG slowing correlates with delirium severity and predicts poor clinical outcomes. Neurology, 2019. 93(13): p. e1260–e1271.

10. Shinozaki, G., et al., Delirium detection by a novel bispectral electroencephalography device in general hospital. Psychiatry Clin Neurosci, 2018. 72(12): p. 856–863.

11. Yamanashi, T., et al., Evaluation of point-of-care thumb-size bispectral electroencephalography device to quantify delirium severity and predict mortality. Br J Psychiatry, 2022. 220(6): p. 322–329.

12. Yamanashi, T., et al., Bispectral EEG (BSEEG) quantifying neuro-inflammation in mice induced by systemic inflammation: A potential mouse model of delirium. J Psychiatr Res, 2021. 133: p. 205–211.

13. Nishiguchi, T., et al., The Bispectral Electroencephalography Method Quantifies Postoperative Delirium-Like States in Young and Aged Male Mice After Head-Mount Implantation Surgery. J Gerontol A Biol Sci Med Sci, 2024. 79(8).

14. Hauglund, N.L., et al., Meningeal Lymphangiogenesis and Enhanced Glymphatic Activity in Mice with Chronically Implanted EEG Electrodes. J Neurosci, 2020. 40(11): p. 2371–2380.

15. Donat, C.K., et al., Microglial Activation in Traumatic Brain Injury. Front Aging Neurosci, 2017. 9: p. 208.

16. Althaus, A.L., et al., Preclinical characterization of zuranolone (SAGE-217), a selective neuroactive steroid GABA(A) receptor positive allosteric modulator. Neuropharmacology, 2020. 181: p. 108333.

17. Patterson, R., et al., Novel neurosteroid therapeutics for post-partum depression: perspectives on clinical trials, program development, active research, and future directions. Neuropsychopharmacology, 2024. 49(1): p. 67–72.

18. Maguire, J.L. and S. Mennerick, Neurosteroids: mechanistic considerations and clinical prospects. Neuropsychopharmacology, 2024. 49(1): p. 73–82.

19. Evered, L., et al., Acute peri-operative neurocognitive disorders: a narrative review. Anaesthesia, 2022. 77 Suppl 1: p. 34–42.

20. Wilson, J.E., et al., Delirium. Nat Rev Dis Primers, 2020. 6(1): p. 90.

21. Kurki, S.N., et al., Acute neuroinflammation leads to disruption of neuronal chloride regulation and consequent hyperexcitability in the dentate gyrus. Cell Rep, 2023. 42(11): p. 113379.

22. Deligiannidis, K.M., et al., Zuranolone for the Treatment of Postpartum Depression. Am J Psychiatry, 2023. 180(9): p. 668–675.

23. Zawilska, J.B. and E. Zwierzyńska, Neuroactive Steroids as Novel Promising Drugs in Therapy of Postpartum Depression-Focus on Zuranolone. Int J Mol Sci, 2025. 26(13).

24. Liu, J., et al., Modeling the efficacy of a novel antidepressant zuranolone for major depressive disorder. J Affect Disord, 2026. 392: p. 120158.

25. Nishiguchi, T., et al., Lipopolysaccharide-Induced Delirium-Like Behavior and Microglial Activation in Mice Correlate With Bispectral Electroencephalography. J Gerontol A Biol Sci Med Sci, 2024. 79(12).

26. Nishiguchi, T., et al., The age-related susceptibility to postoperative delirium quantified by bispectral electroencephalography correlates with postoperative delirium-like behavior in mice. PCN Rep, 2025. 4(3): p. e70142.

27. Nishiguchi, T., et al., Discovery of novel protective agents for infection-related delirium through bispectral electroencephalography. Transl Psychiatry, 2024. 14(1): p. 413.

28. Nishizawa, Y., et al., Bispectral EEG (BSEEG) Algorithm Captures High Mortality Risk Among 1,077 Patients: Its Relationship to Delirium Motor Subtype. Am J Geriatr Psychiatry, 2023. 31(9): p. 704–715.

29. Shimura, A., et al., A web-based delirium detection application using bispectral electroencephalography (BSEEG). Gen Hosp Psychiatry, 2024. 91: p. 223–224.

30. Aldecoa, C., et al., Update of the European Society of Anaesthesiology and Intensive Care Medicine evidence-based and consensus-based guideline on postoperative delirium in adult patients. Eur J Anaesthesiol, 2024. 41(2): p. 81–108.

31. Tateda, K., et al., Lipopolysaccharide-induced lethality and cytokine production in aged mice. Infect Immun, 1996. 64(3): p. 769–74.

32. Aoyama, B., et al., Role of neurosteroid allopregnanolone on age-related differences in exercise-induced hypoalgesia in rats. J Pharmacol Sci, 2019. 139(2): p. 77–83.

33. Kozai, T.D., et al., In vivo two-photon microscopy reveals immediate microglial reaction to implantation of microelectrode through extension of processes. J Neural Eng, 2012. 9(6): p. 066001.

34. Kozai, T.D., et al., Brain tissue responses to neural implants impact signal sensitivity and intervention strategies. ACS Chem Neurosci, 2015. 6(1): p. 48–67.

35. Spagnoli, G., et al., Glial Response and Neuronal Modulation Induced by Epidural Electrode Implant in the Pilocarpine Mouse Model of Epilepsy. Biomolecules, 2024. 14(7).

