## Supplementary Material for "Zuranolone mitigates delirium-like bispectral EEG changes, behavioral deficits, and neuroinflammation across surgical and inflammatory mouse models and age groups"

### Supplementary Methods

#### Experimental design and overview

This study employed two complementary murine models of delirium-like brain dysfunction: a postoperative delirium (POD) model induced by surgical stress associated with EEG head-mount implantation, and a systemic inflammation model induced by lipopolysaccharide (LPS) administration. Experiments were conducted across young, aged, and super-aged cohorts to assess age-dependent susceptibility and treatment response. Zuranolone or vehicle was administered using either prevention or treatment paradigms, depending on the experiment.

---

#### Zuranolone formulation, dosing, and administration

Zuranolone (>98% by HPLC) was supplied as a powder and freshly dissolved in 30% (w/v) sulfobutylether-β-cyclodextrin (SBE-β-CD) immediately before administration. Vehicle-treated animals received an equivalent volume of 30% SBE-β-CD alone.

Zuranolone or vehicle was administered by oral gavage at the following doses:

- **Young POD model:** Zuranolone was administered at 1 mg $kg^{-1}$ in all experiments.
- **Young LPS model:**
    - BSEEG experiments: Zuranolone was administered at 0.5 or 1 mg $kg^{-1}$.
    - Behavioral testing and immunofluorescence experiments: Zuranolone was administered at 1 mg $kg^{-1}$.
- **Aged and super-aged cohorts:**
    - POD model: Zuranolone was administered at 1 mg $kg^{-1}$.
    - LPS model: Zuranolone was administered at 0.5 mg $kg^{-1}$.

All doses were selected based on prior pharmacological validation and pilot experiments using the same experimental platforms.

---

#### Detailed experimental paradigms

##### Postoperative delirium (POD) model

In the POD model, the surgical procedure required for EEG head–mount implantation served as the delirium–inducing insult. Behavioral testing was performed one day before surgery to obtain baseline (pre–test) values. All mice then underwent surgery and were administered either vehicle or Zuranolone according to group allocation. Behavioral post–testing was conducted 24 h after surgery. Accordingly, behavioral comparisons reflect surgery + vehicle versus surgery + Zuranolone.

EEG recording commenced immediately after surgery and continued continuously for the duration of the experiment.

---

##### LPS–induced systemic inflammation model

Systemic inflammation was induced by intraperitoneal injection of LPS (*Escherichia coli* O111:B4). Behavioral baseline testing was performed 24 h before LPS administration. In the prevention/early–intervention paradigm, vehicle or zuranolone was administered immediately before LPS injection; in the treatment paradigm, dosing was initiated after LPS injection as specified in Supplementary Table 1. Behavioral post–testing was conducted 24 h after LPS injection. Thus, behavioral comparisons reflect LPS + vehicle versus LPS + Zuranolone.

EEG recordings were conducted continuously before and after LPS administration to capture dynamic changes in brain electrical activity.

---

#### Behavioral testing procedures

##### Buried food test (BFT)

The BFT was performed as previously described with minor modifications [25,26]. Twenty-four hours before testing, all chow pellets were removed from the home cage while water remained available ad libitum. A clean test cage was prepared with fresh bedding (3 cm depth), and a ~2 g food pellet identical to the home-cage chow was buried 0.5 cm below the bedding surface at a randomized location for each mouse. Mice were placed in the center of the cage, and latency was defined as the time until the mouse uncovered the pellet and grasped it with forepaws and teeth. The maximum observation time was 5 min; mice not retrieving the pellet were assigned a latency of 300 s. A new cage was used for each mouse, and gloves were changed between animals.

##### Open-field test (OFT)

OFT was performed as previously described [25,26]. Mice were placed in the center of an open-field arena (40 × 40 × 40 cm) and allowed to explore freely for 5 min. Total distance traveled, time spent in the center zone, freezing time, and latency to first center entry were quantified by video tracking (SMART 3.0). The arena was cleaned with 70% ethanol between animals.

##### Y-maze test

The Y-maze test was performed as previously described [25,26]. The Y-maze consisted of three arms (5 × 35 × 10 cm; 120° between arms) and was conducted as a two-trial novel-arm recognition task. During the 10-min training trial, the novel arm was blocked and mice explored the start arm and one familiar arm. After a 2-h inter-trial interval, the 5-min retention trial was performed with free access to all three arms. Time spent in and entries into the novel arm were quantified using video tracking (SMART 3.0). The maze was cleaned with 70% ethanol between trials.

All behavioral testing was conducted during the light phase under standardized environmental conditions. Apparatuses were cleaned between animals to minimize olfactory cues.

#### Composite Z score calculation (expanded description)

To integrate multimodal behavioral outcomes into a single quantitative index of delirium-like severity, baseline-corrected behavioral measures were combined into a composite Z score using a standardized algorithm applied consistently across experimental models.

##### Baseline correction

For each animal and each behavioral variable, baseline correction was calculated as:

$$\Delta X = X_{\text{post}} - X_{\text{pre}}$$

where $X_{\text{pre}}$ represents the baseline value obtained during pre-testing and $X_{\text{post}}$ represents the value obtained during post-testing (24 h after surgery or 24 h after LPS administration, depending on the model).

For behavioral measures in which lower values indicate better performance (e.g., latency to eat in the buried food test), values were sign-inverted so that higher values consistently represented greater behavioral impairment.

##### Z score calculation

For each behavioral variable, an individual Z score was calculated as:

$$Z = \frac{\Delta X - \mu_{\text{ref}}}{\sigma_{\text{ref}}}$$

where $\mu_{\text{ref}}$ and $\sigma_{\text{ref}}$ represent the mean and standard deviation of the reference distribution derived from the corresponding vehicle-treated control group within the same experimental model.

##### Reference distributions

Behavioral experiments were conducted exclusively in young mice. Reference distributions were predefined prior to statistical analysis to avoid circularity and were applied uniformly across treatment groups within each experimental model.

- **POD model:** Reference distributions were derived from baseline-corrected behavioral changes (24 h post – pre) in mice that underwent surgery and received vehicle administration.
- **LPS model:** Reference distributions were derived from baseline-corrected behavioral changes (24 h post – pre) in vehicle-treated mice subjected to LPS challenge.

**Composite score**

Individual Z scores from the following six behavioral parameters were aggregated to generate a composite Z score per mouse:

1. Latency to eat food (BFT)
2. Time spent in the center (OFT)
3. Latency to center entry (OFT)
4. Freezing time (OFT)
5. Entries into the novel arm (Y-maze)
6. Time spent in the novel arm (Y-maze)

The composite Z score was calculated as the sum of individual Z scores and normalized by the standard deviation of this sum in the reference group. This normalization ensured comparability of composite Z score distributions across models. This composite metric served as an integrated quantitative index of delirium-like behavioral severity analogous to clinically used delirium severity scales.

**Supplementary Table 1 | Group sizes, dosing regimens, and administration schedules for the POD and LPS models**

| Model | Age | Paradigm | Outcome | Group | n | Z dose (mg kg$^{-1}$) | Z route | Z timing (relative to insult) | LPS dose (mg kg$^{-1}$) | Route |
|---|---|---|---|---|---|---|---|---|---|---|
| POD | Young | Prevention | BSEEG | Vehicle | 11 | 0 (vehicle) | Oral | 10 min before surgery | - | - |
| POD | Young | Prevention | BSEEG | Z | 10 | 1.0 | Oral | 10 min before surgery | - | - |
| POD | Young | Treatment | BSEEG | Vehicle | 8 | 0 (vehicle) | Oral | 24 $\pm$ 1 h after surgery | - | - |
| POD | Young | Treatment | BSEEG | Z | 9 | 1.0 | Oral | 24 $\pm$ 1 h after surgery | - | - |
| LPS | Young | Prevention | BSEEG | Vehicle | 9 | 0 (vehicle) | Oral | Immediately before LPS | 1.0 | i.p. |
| LPS | Young | Prevention | BSEEG | Z | 8 | 0.5 | Oral | Immediately before LPS | 1.0 | i.p. |
| LPS | Young | Prevention | BSEEG | Z | 8 | 1.0 | Oral | Immediately before LPS | 1.0 | i.p. |
| LPS | Young | Treatment | BSEEG | Vehicle | 9 | 0 (vehicle) | Oral | 6 h after LPS administration | 1.0 | i.p. |
| LPS | Young | Treatment | BSEEG | Z | 8 | 1.0 | Oral | 6 h after LPS administration | 1.0 | i.p. |
| POD | Aged | Prevention | BSEEG | Vehicle | 6 | 0 (vehicle) | Oral | 10 min before surgery | - | - |
| POD | Aged | Prevention | BSEEG | Z | 7 | 1.0 | Oral | 10 min before surgery | - | - |
| LPS | Aged | Prevention | BSEEG | Vehicle | 7 | 0 (vehicle) | Oral | Immediately before LPS | 0.1 | i.p. |
| LPS | Aged | Prevention | BSEEG | Z | 7 | 0.5 | Oral | Immediately before LPS | 0.1 | i.p. |
| POD | Super Aged | Prevention | BSEEG +Survival | Vehicle | 10 | 0 (vehicle) | Oral | 10 min before surgery | - | - |
| POD | Super Aged | Prevention | BSEEG +Survival | Z | 9 | 1.0 | Oral | 10 min before surgery | - | - |
| LPS | Super Aged | Prevention | BSEEG +Survival | Vehicle | 11 | 0 (vehicle) | Oral | Immediately before LPS | 0.05 | i.p. |
| LPS | Super Aged | Prevention | BSEEG +Survival | Z | 11 | 0.5 | Oral | Immediately before LPS | 0.05 | i.p. |

- **Zuranolone = Z, intraperitoneal = i.p.**
- **Administration immediately before LPS injection was defined a priori as a prevention/early-intervention paradigm, as it was performed before the onset of delirium-like behavioral and electrophysiological abnormalities in this model.**

**Supplementary Fig. 1 | sBSEEG scores for each prevention group in the young-mouse postoperative delirium (POD) model**

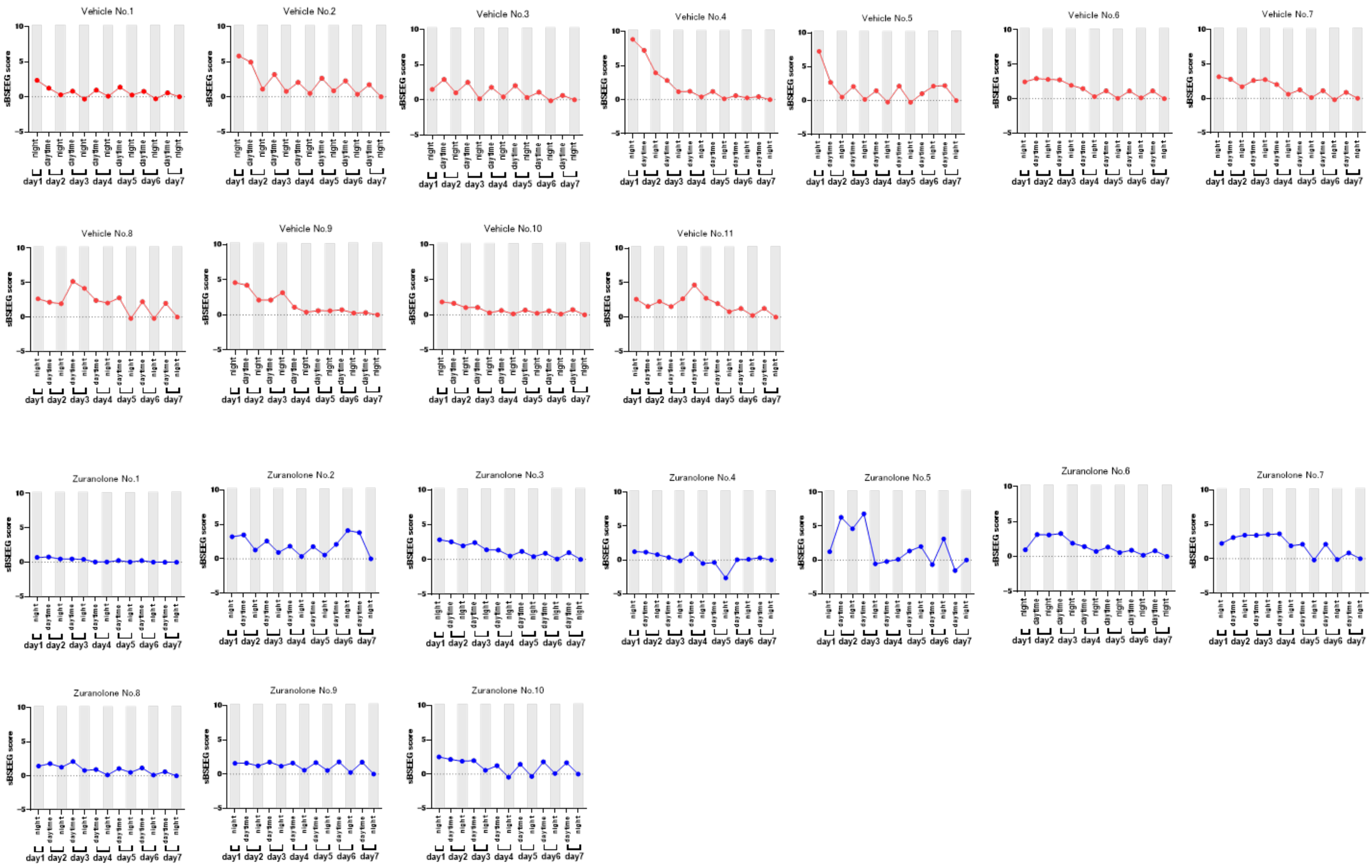

**Supplementary Fig. 2 | sBSEEG scores for each treatment group in the young-mouse POD model**

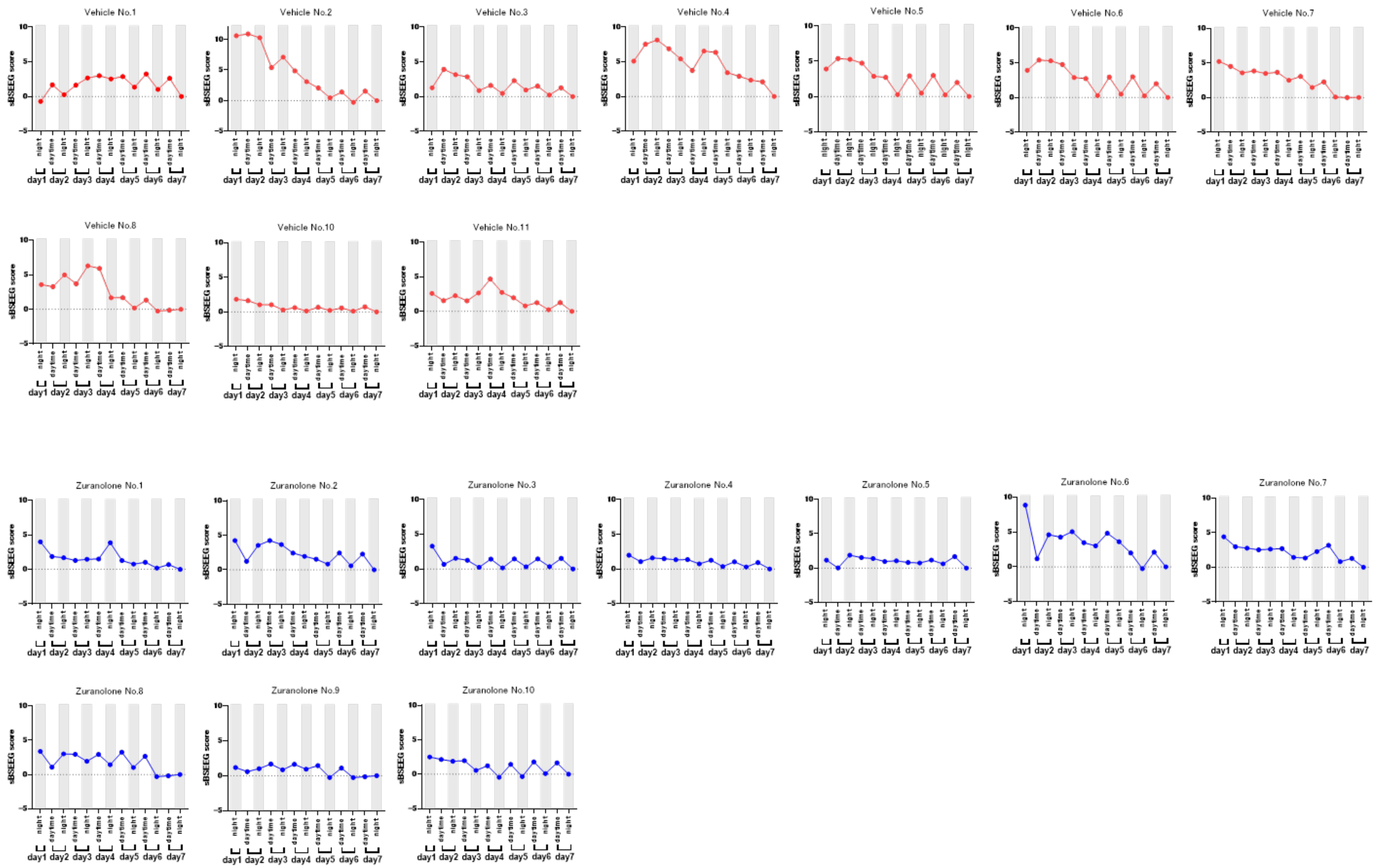

**Supplementary Fig. 3 | sBSEEG scores for each prevention group in the young-mouse lipopolysaccharide (LPS) model**

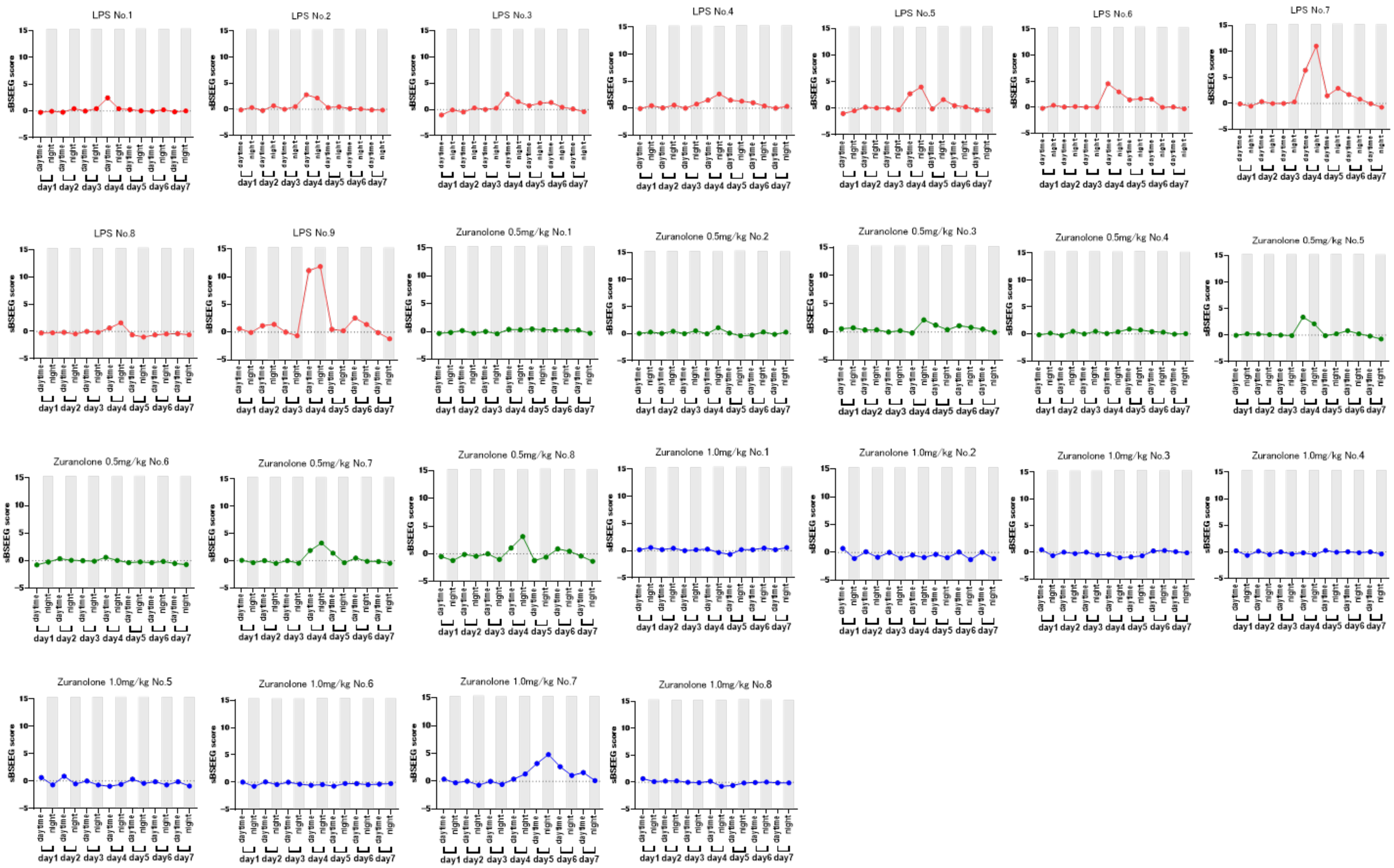

**Supplementary Fig. 4 | sBSEEG scores for each treatment group in the young-mouse LPS model**

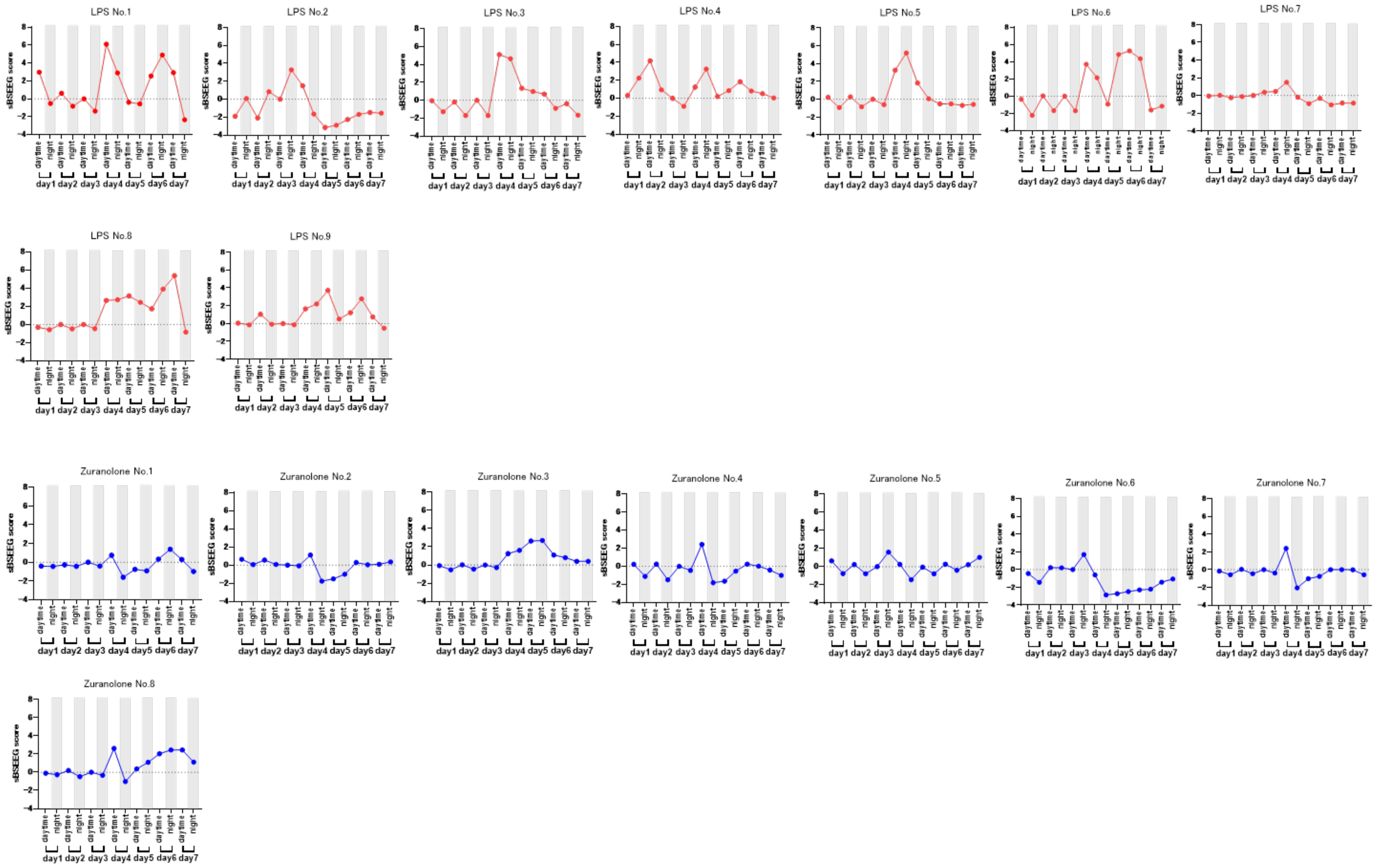

**Supplementary Fig. 5 | sBSEEG scores and experimental timeline for combined prevention and treatment groups in the young-mouse LPS model**

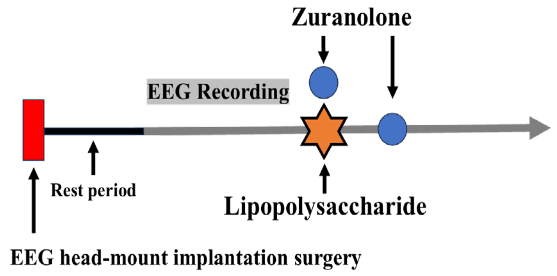


**Young-mouse LPS Model - Combined Prevention and Treatment**

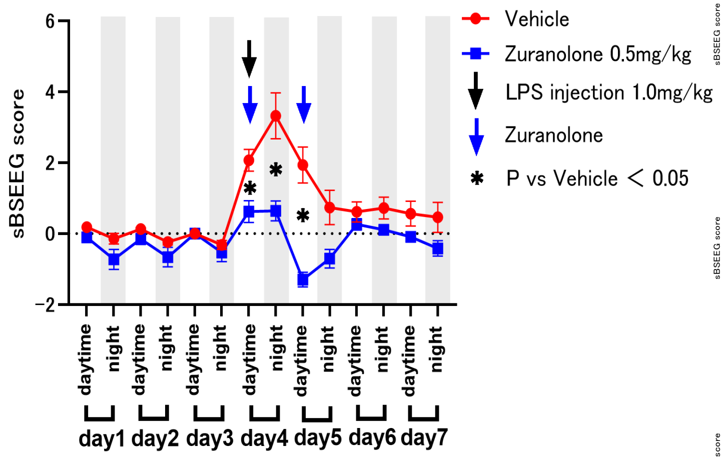


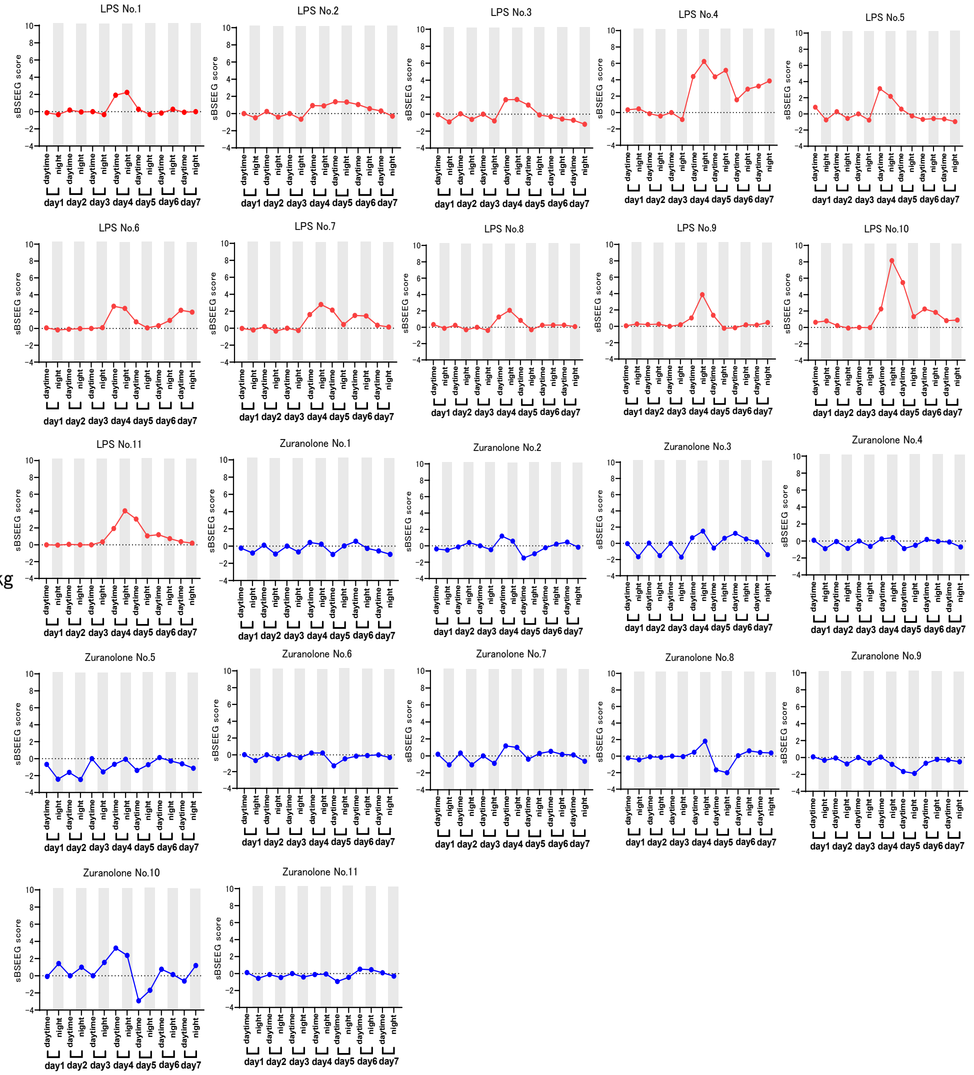


**In a separate cohort, zuranolone or vehicle was administered once daily for two consecutive days: the first dose was given immediately before LPS injection (prevention/early-intervention), and the second dose was given 24 h after LPS injection (treatment).**

**Supplementary Fig. 6 | sBSEEG scores and experimental timeline for prevention in young female mice in the LPS model**

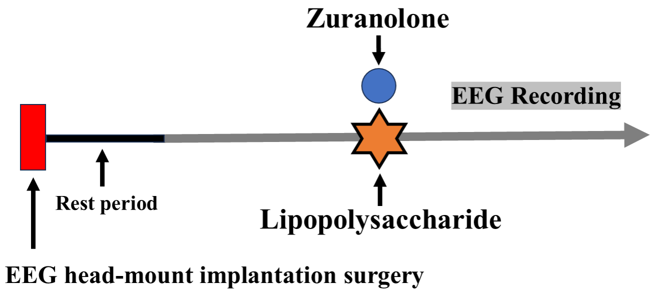


**Female young-mouse LPS Model - Prevention**

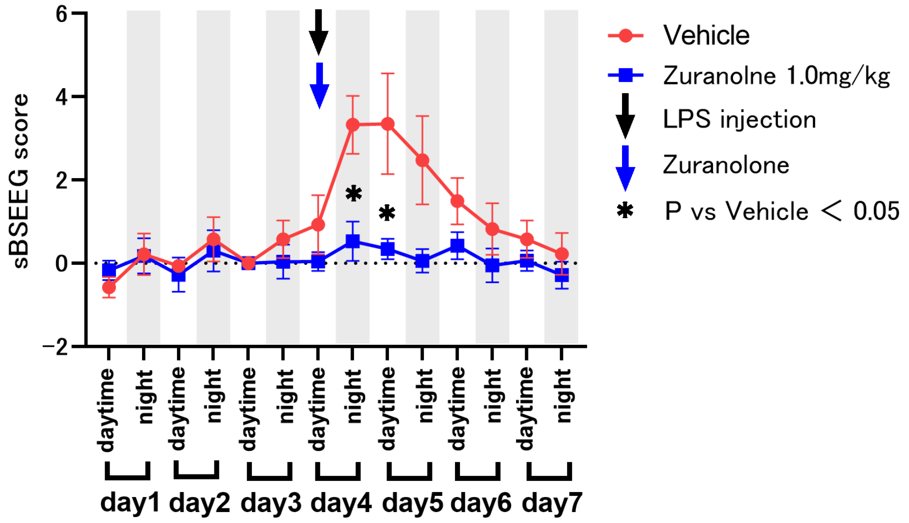


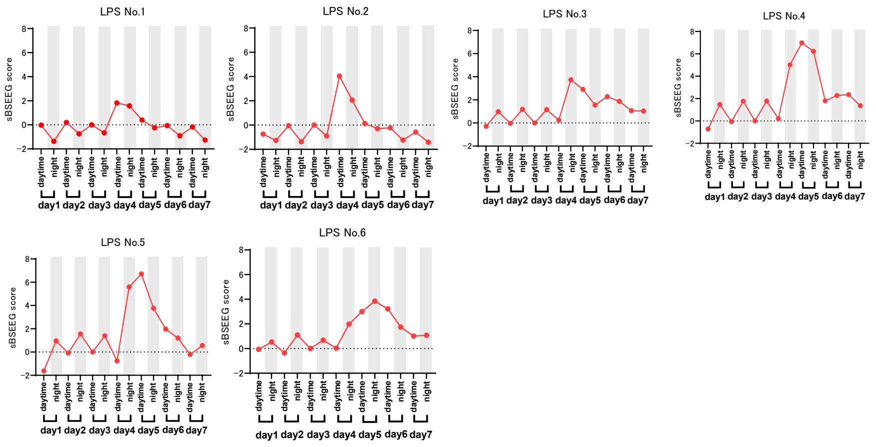


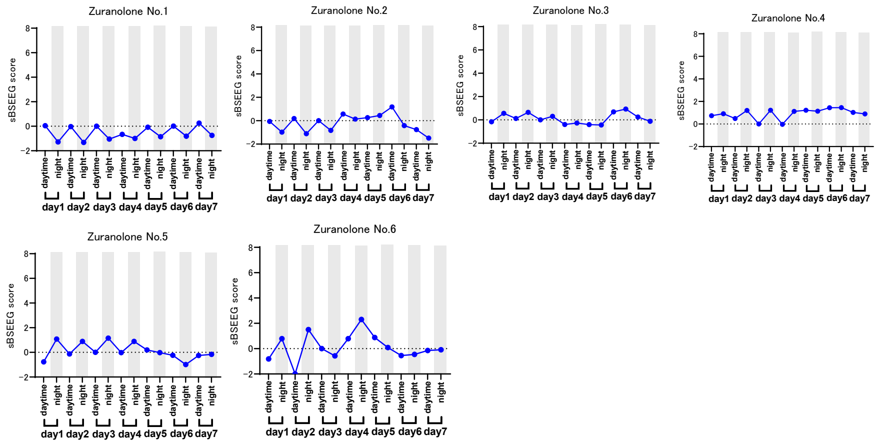

**Supplementary Fig. 7 | sBSEEG scores following SBE-β-CD and zuranolone administration in young male mice**

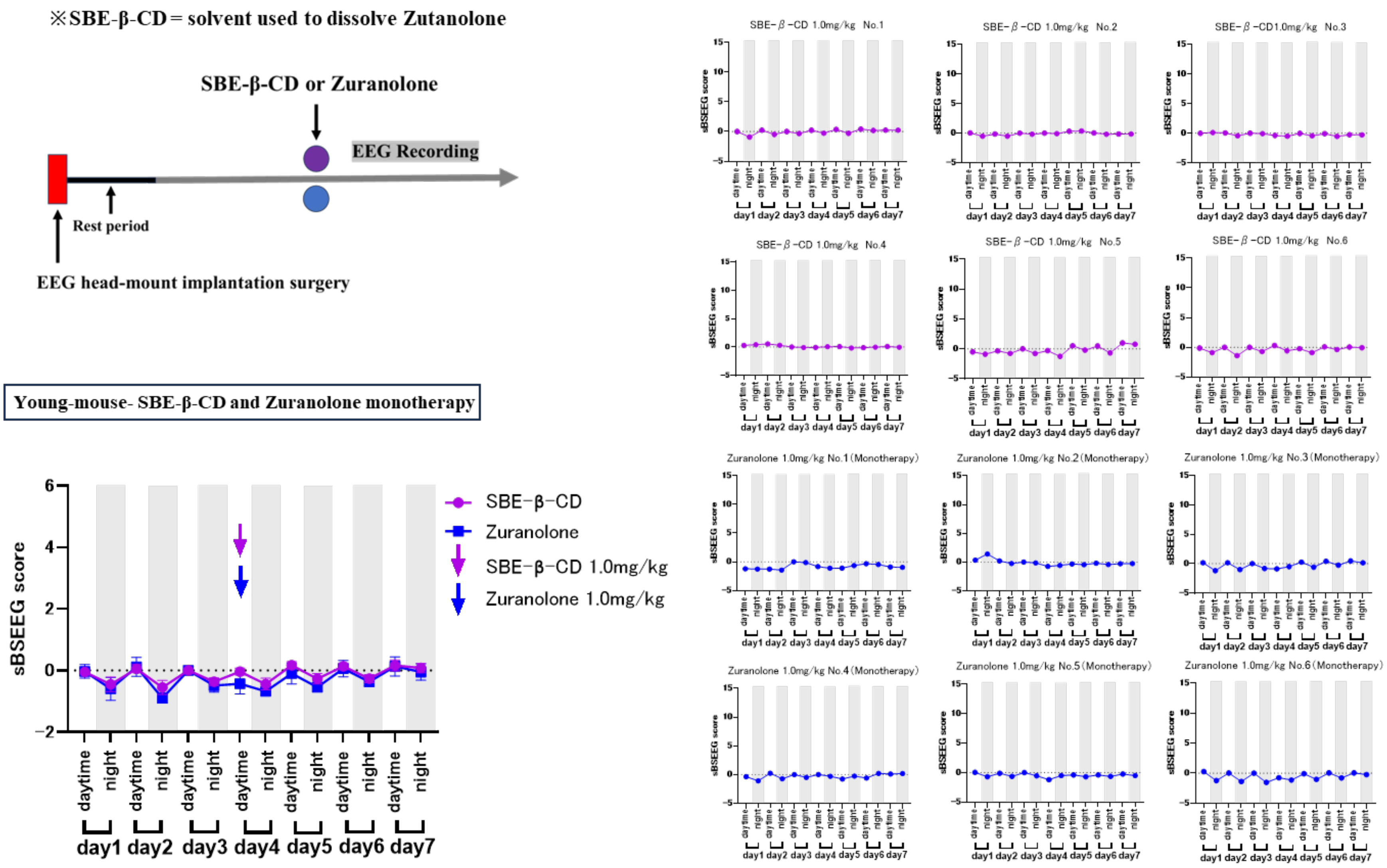

### Supplementary Fig. 8 | sBSEEG scores for each prevention group in the aged-mouse POD model

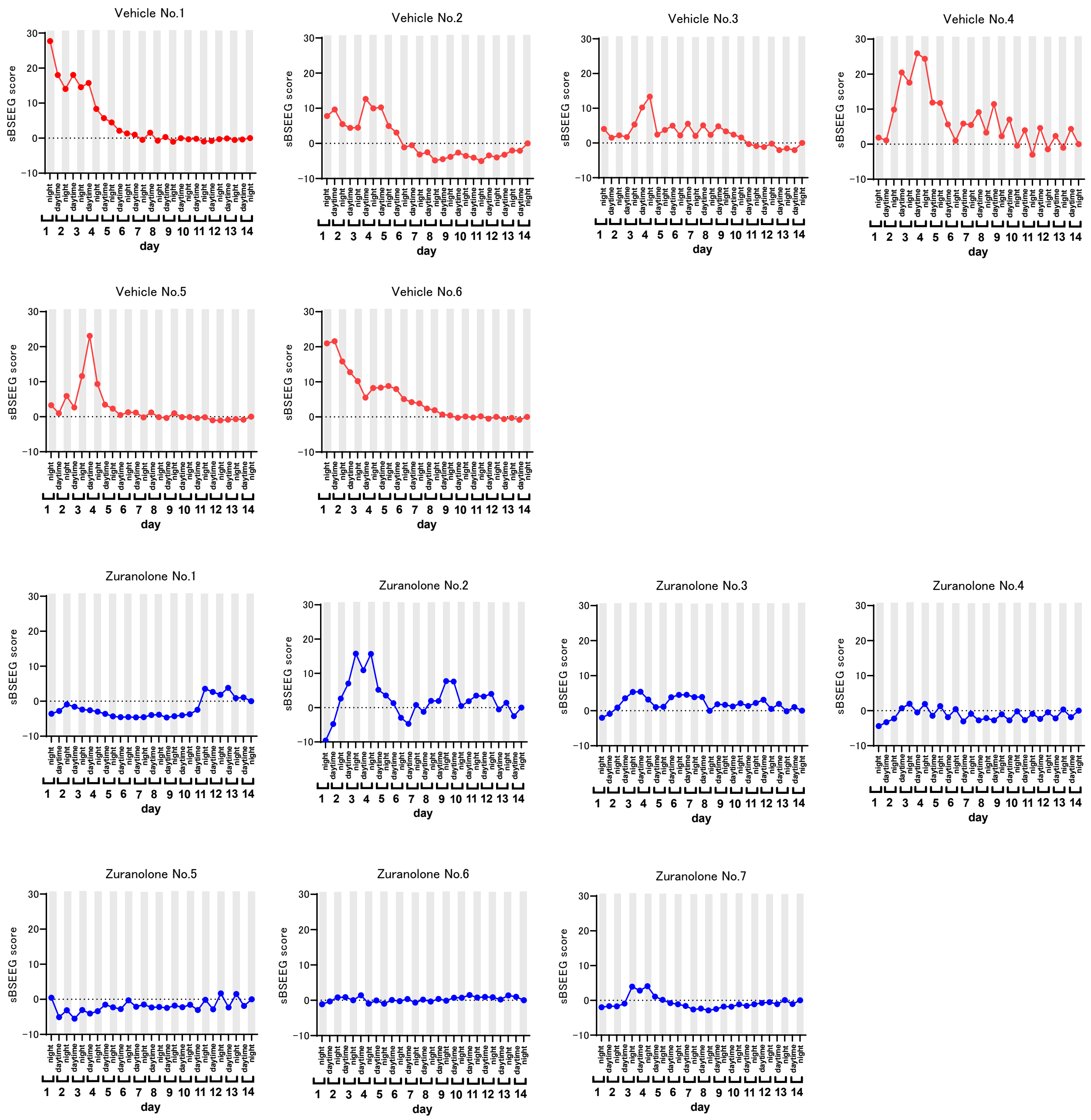

**Supplementary Fig. 9 | sBSEEG scores for each prevention group in the aged-mouse lipopolysaccharide (LPS) model**

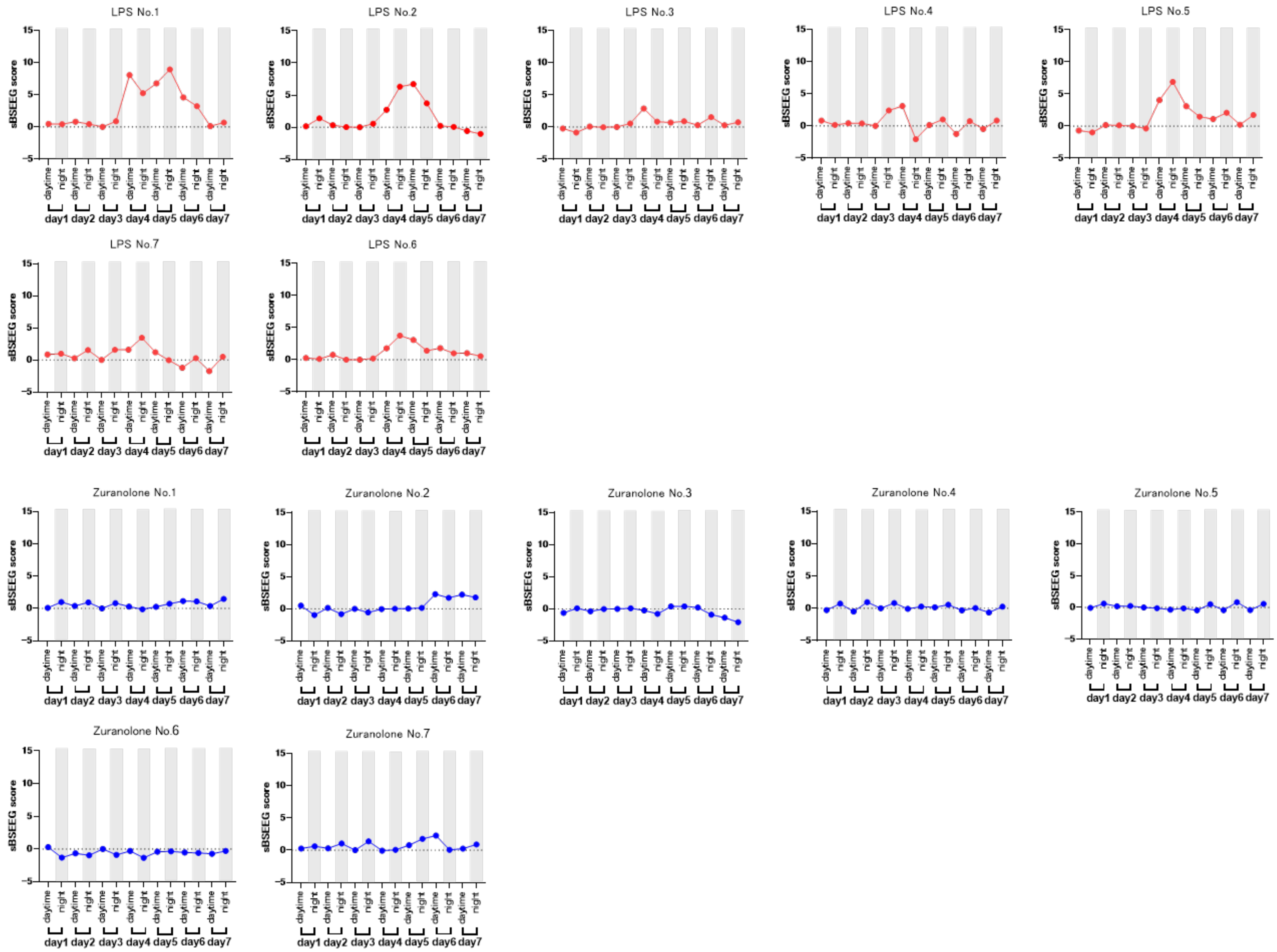

**Supplementary Fig. 10 | sBSEEG scores and experimental timeline for prevention in the super-aged mouse POD model**

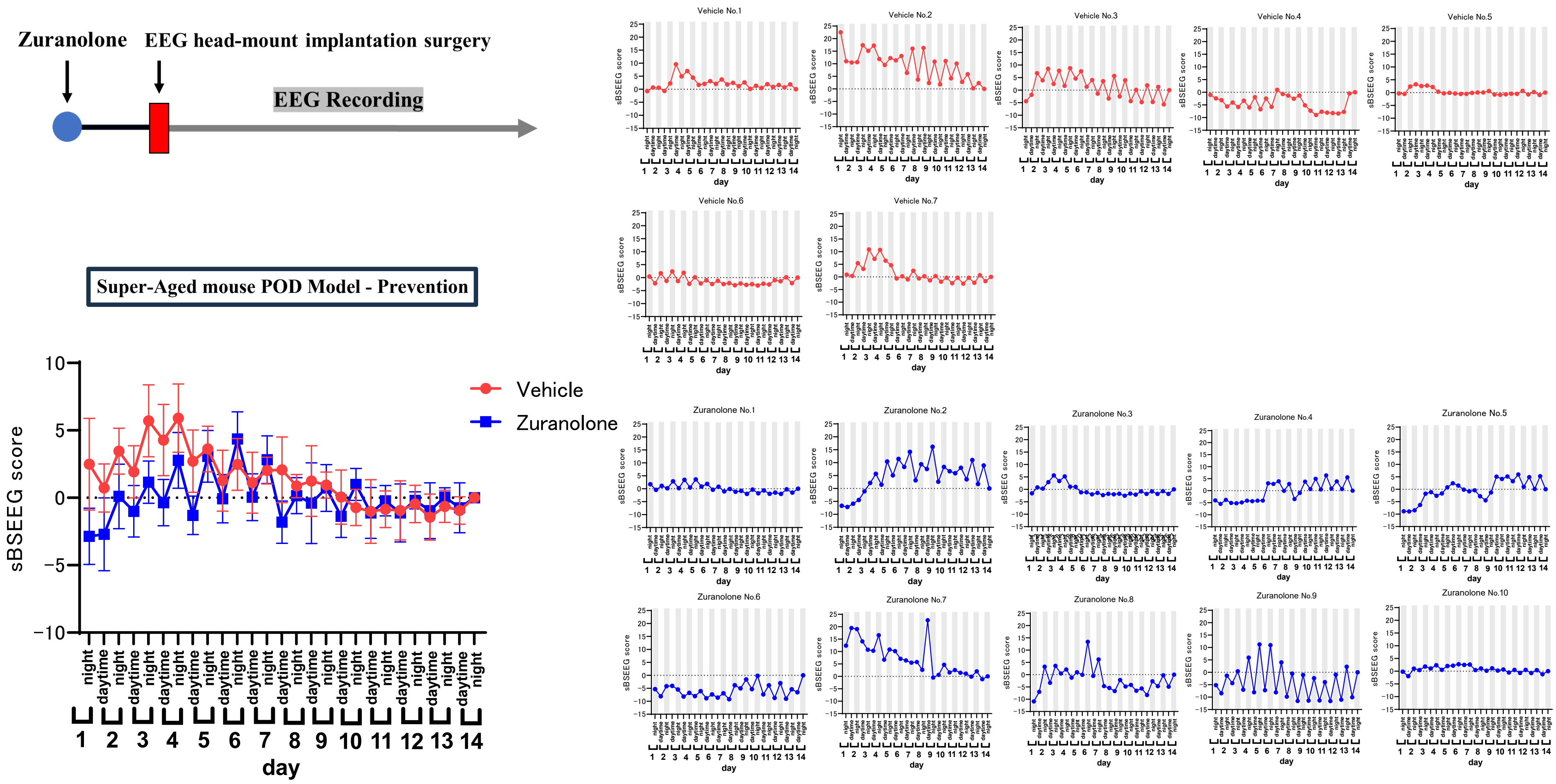

**Supplementary Fig. 11 | sBSEEG scores and experimental timeline for prevention in the super-aged mouse LPS model**

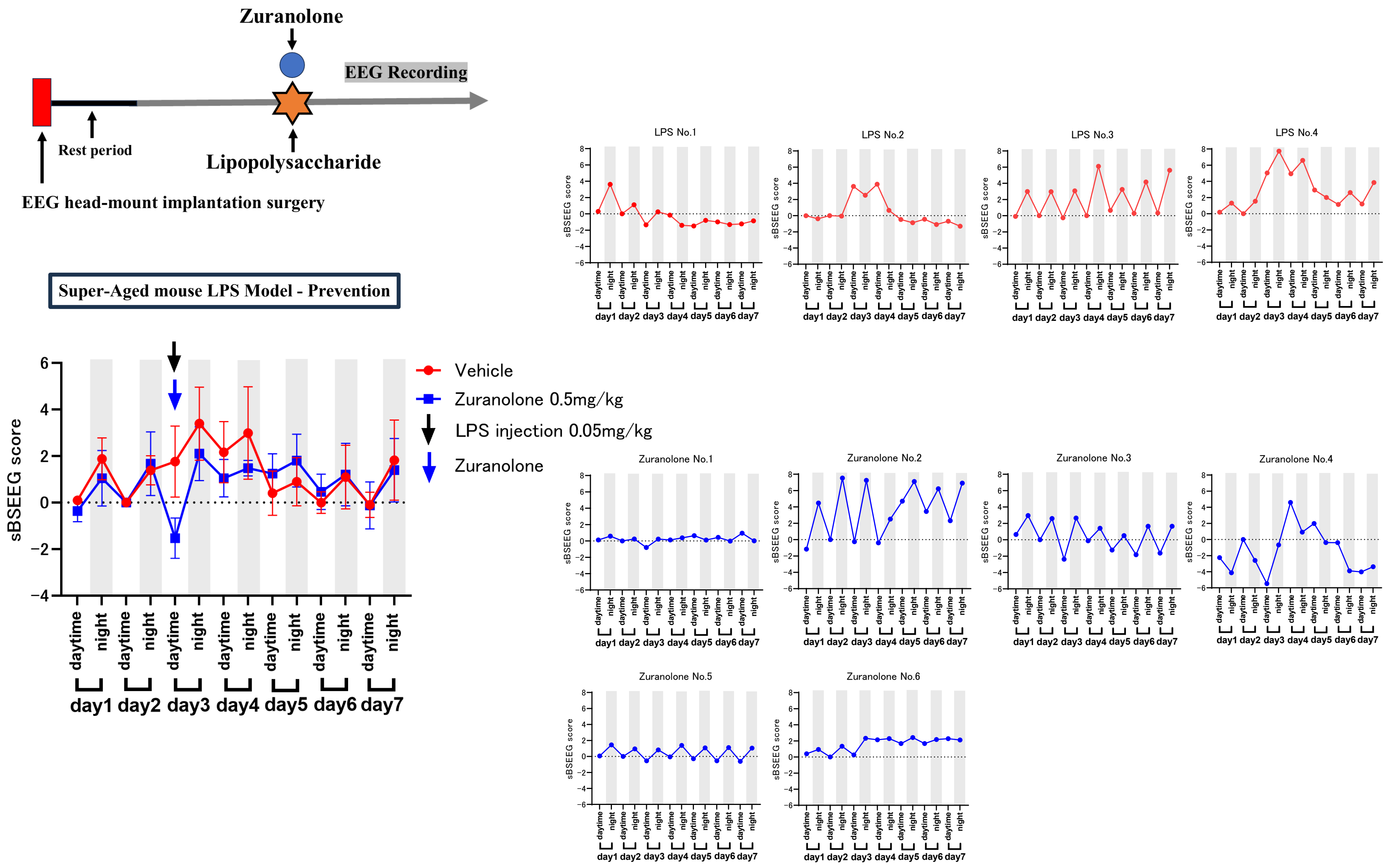

**Supplementary Table 2 | Z scores of young mice POD model**

| Vehicle | No.1 | No.2 | No.3 | No.4 | No.5 | No.6 | No.7 | No.8 | No.9 | No.10 | No.11 | No.12 | No.13 |
|---|---|---|---|---|---|---|---|---|---|---|---|---|---|
| **BFT: Latency to pellet** | -1.28274 | -0.66933 | 0.80625 | -0.26265 | 0.285143 | -0.21917 | -0.30678 | -0.79027 | -0.42499 | -0.64069 | 0.659278 | 2.883131 | -0.03718 |
| **OFT: Latency to center** | -1.09394 | -0.16707 | -0.76716 | -0.29593 | 3.11029 | -0.23873 | -0.25888 | 0.184569 | -0.47272 | -0.0062 | -0.24198 | -0.56697 | 0.814724 |
| **OFT: Freezing time** | 1.631518 | 1.346475 | 0.681527 | 1.094941 | -0.19601 | 0.229626 | -0.61418 | 0.449464 | -0.23812 | -2.00113 | -0.54649 | -0.65131 | -1.1863 |
| **OFT: Time spent in center** | -0.26634 | -0.2454 | -0.60043 | 0.401832 | 0.65165 | 0.206365 | -1.52242 | -0.83729 | -0.05991 | -1.65805 | 0.883517 | 1.02912 | -0.26285 |
| **YMZ: Entries in novel arm** | 2.260701 | -1.25594 | 0.033492 | -0.90428 | 0.385156 | 0.150713 | 0.502378 | 0.267935 | -1.0215 | -1.37317 | -0.55262 | 0.385156 | -0.43539 |
| **YMZ: Duration in novel arm** | -0.69516 | -2.07609 | 1.456048 | -0.34604 | 0.015532 | -0.04184 | -0.39096 | 1.960384 | -0.60977 | 0.710217 | -0.28333 | 0.845863 | -0.54484 |
| **Composite Z score** | 0.212441 | -1.17616 | 0.617239 | -0.11968 | 1.630312 | 0.033345 | -0.99344 | 0.473476 | -1.084 | -1.90534 | -0.0313 | 1.505015 | -0.63339 |

| Zuranolone | No.1 | No.2 | No.3 | No.4 | No.5 | No.6 | No.7 | No.8 | No.9 | No.10 | No.11 | No.12 | No.13 |
|---|---|---|---|---|---|---|---|---|---|---|---|---|---|
| **BFT: Latency to pellet** | -3.35962 | -0.18676 | -1.28404 | -1.36566 | -1.12769 | -0.2525 | -1.28078 | -1.52721 | -0.43983 | -0.37343 | -0.29481 | 0.683622 | -0.41848 |
| **OFT: Latency to center** | -0.43275 | -0.77204 | -0.7475 | -0.66625 | -0.18689 | -0.56421 | -0.61881 | -0.54845 | -0.40236 | -0.23386 | 0.253142 | -0.15407 | 0.0029 |
| **OFT: Freezing time** | 1.576275 | -0.95605 | 0.156044 | 1.661856 | 0.017712 | 1.132524 | -1.29611 | 0.347582 | 1.099242 | -0.84557 | -0.68618 | 0.376562 | -0.51049 |
| **OFT: Time spent in center** | 0.014888 | 0.592312 | -1.98216 | 0.223818 | -0.52165 | 0.764841 | -0.3202 | -0.20651 | 0.033836 | 0.712982 | 1.47241 | 2.488137 | 0.954822 |
| **YMZ: Entries in novel arm** | -0.31817 | -0.20095 | -0.20095 | -0.31817 | -0.31817 | -1.0215 | -0.20095 | -0.20095 | 0.033492 | 0.502378 | 0.736821 | 0.385156 | -0.08373 |
| **YMZ: Duration in novel arm** | -3.96401 | -0.40519 | 1.092694 | 1.488514 | 0.306392 | 0.077795 | -1.80746 | 0.600366 | 0.655514 | 1.552112 | 1.535212 | -1.21907 | 0.574571 |
| **Composite Z score** | -2.48602 | -0.73954 | -1.13726 | 0.392685 | -0.70182 | 0.052514 | -2.11827 | -0.58865 | 0.375736 | 0.504083 | 1.156699 | 0.981747 | 0.199237 |

Individual behavioral Z scores and composite Z scores are shown for each animal. Behavioral values were baseline-corrected, sign-adjusted so that higher values indicate greater impairment, and normalized using model-specific reference distributions as described in the Methods and Supplementary Methods. Composite Z scores represent the aggregated behavioral impairment for each mouse.
No missing behavioral values were present for animals included in this table.

**Supplementary Table 3 | Z scores of young mice LPS model**

| Vehicle | No.1 | No.2 | No.3 | No.4 | No.5 | No.6 | No.7 | No.8 | No.9 | No.10 | No.11 | No.12 | No.13 |
|---|---|---|---|---|---|---|---|---|---|---|---|---|---|
| **BFT: Latency to pellet** | 0.3828287 | -1.986959 | 0.8856078 | -0.66302 | 0.8530983 | 0.5006093 | 1.3654704 | 0.6019752 | 0.889978 | -1.113464 | 0.1955523 | -1.714838 | 0.373875 |
| **OFT: Latency to center** | -1.070297 | 1.0806295 | 0.6782278 | 1.7889943 | -0.235643 | -0.714392 | -1.910117 | -0.879716 | 1.1724758 | 0.1265757 | 0.3613579 | 0.6311565 | 0.027841 |
| **OFT: Freezing time** | -0.342665 | 2.492897 | -1.699806 | 0.4017862 | -0.188744 | -0.815208 | -0.365424 | -0.436395 | -1.412626 | 0.280506 | 0.5784661 | 0.8024602 | -0.02973 |
| **OFT: Time spent in center** | 1.3609916 | 1.0525922 | -1.087626 | 0.3461531 | -0.004047 | 2.0056579 | 0.2620864 | -1.656121 | -1.21628 | 0.2592997 | -0.710486 | 0.2955273 | -0.03424 |
| **YMZ: Entries in novel arm** | 1.0582498 | -0.439947 | 2.0570473 | 0.0594522 | -0.606413 | 0.558851 | 0.0594522 | -0.939345 | 1.224716 | 0.2259185 | -1.105812 | -1.60521 | 0.558851 |
| **YMZ: Duration in novel arm** | 1.395869 | 0.5252011 | 0.09611 | -1.67977 | 0.2317877 | -0.038735 | -1.398009 | 0.2334524 | 1.8207983 | -0.049972 | -0.132794 | 0.041173 | 0.701665 |
| **Composite Z score** | 1.550252 | 1.265017 | 0.43162 | 0.117751 | 0.023234 | 0.694995 | -0.9224 | -1.42834 | 1.151094 | -0.1259 | -0.37783 | -0.71958 | 0.742116 |

| Zuranolone | No.1 | No.2 | No.3 | No.4 | No.5 | No.6 | No.7 | No.8 | No.9 | No.10 | No.11 | No.12 | No.13 |
|---|---|---|---|---|---|---|---|---|---|---|---|---|---|
| **BFT: Latency to pellet** | -2.082995 | -1.715477 | -0.885364 | -0.671974 | -0.693398 | -1.018387 | -1.682115 | 0.1545681 | -1.454015 | -1.385798 | -0.003342 | -0.975858 | -0.3666 |
| **OFT: Latency to center** | -1.28384 | 0.4652591 | -0.07893 | -1.304505 | -2.586335 | -1.796457 | -1.066853 | -0.670625 | -3.202854 | -0.237939 | -0.187424 | -1.595543 | -0.75285 |
| **OFT: Freezing time** | -1.610867 | -0.475624 | 1.7397617 | 0.3907063 | 0.8123423 | -0.265105 | -0.889174 | 0.1023506 | 0.5284567 | 0.9794395 | 0.8225238 | 1.2315826 | -0.05998 |
| **OFT: Time spent in center** | 0.2407214 | 0.1924179 | 0.1831287 | -0.756003 | 0.1938113 | -0.825672 | -0.030521 | -0.896633 | 0.2235365 | -1.680273 | -0.906023 | 0.1064933 | -0.12852 |
| **YMZ: Entries in novel arm** | 0.558851 | -0.107014 | -0.772879 | -0.606413 | -0.772879 | -0.107014 | 0.8917835 | 0.1054093 | -0.27348 | -0.772879 | 0.8917835 | -1.272278 | 1.557649 |
| **YMZ: Duration in novel arm** | 1.807064 | 0.2738228 | 1.0737386 | 0.3000427 | 0.3595578 | 0.1918335 | 1.3554987 | -0.118297 | 1.0849757 | 0.1989087 | 0.4311423 | 1.8220469 | 0.775331 |
| **Composite Z score** | -1.10095 | -0.63456 | 0.584798 | -1.2296 | -1.2476 | -1.7741 | -0.65998 | -0.61441 | -1.43634 | -1.34587 | 0.486921 | -0.31739 | 0.475949 |

Individual behavioral Z scores and composite Z scores are shown for each animal. Behavioral values were baseline-corrected, sign-adjusted so that higher values indicate greater impairment, and normalized using model-specific reference distributions as described in the Methods and Supplementary Methods. Composite Z scores represent the aggregated behavioral impairment for each mouse.
No missing behavioral values were present for animals included in this table.
